# Homeostasis at different backgrounds: The roles of overlayed feedback structures in vertebrate photoadaptation

**DOI:** 10.1101/2023.01.25.525568

**Authors:** Jonas V. Grini, Melissa Nygård, Peter Ruoff

## Abstract

We have studied the resetting behavior of eight basic integral controller motifs with respect to different but constant backgrounds. We found that the controllers split symmetrically into two classes: one class, based on derepression of the compensatory flux, leads to more rapid resetting kinetics as backgrounds increase. The other class, which directly activates the compensatory flux, shows a slowing down in the resetting at increased backgrounds. We found a striking analogy between the resetting kinetics of vertebrate photoreceptors and controllers based on derepression, i.e. vertebrate rod or cone cells show decreased sensitivities and accelerated response kinetics as background illuminations increase. The central molecular model of vertebrate photoadaptation consists of an overlay of three negative feedback loops with cytosolic calcium 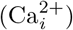, cyclic guanosine monophosphate (cGMP) and cyclic nucleotide-gated (CNG) channels as components. While in one of the feedback loops the extrusion of 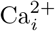 by potassium-dependent sodium-calcium exchangers (NCKX) can lead to integral control with cGMP as the controlled variable, the expected robust perfect adaptation of cGMP is lost, because of the two other feedback loops. They avoid that 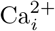 levels become too high and toxic. Looking at psychophysical laws, we found that in all of the above mentioned basic controllers Weber’s law is followed when a “just noticeable difference” (threshold) of 1% of the controlled variable’s set-point was considered. Applying comparable threshold pulses or steps to the photoadaptation model we find, in agreement with experimental results, that Weber’s law is followed for relatively high backgrounds, while Stephens’ power law gives a better description when backgrounds are low. Limitations of our photoadaption model, in particular with respect to potassium/sodium homeostasis, are discussed. Finally, we discuss possible implication of background perturbations in biological controllers when compensatory fluxes are based on activation.

## Introduction

In 1929 Walter B. Cannon [1] defined homeostasis as the sum of the physiological processes which keep the steady states in a cell or organism within narrow limits [2]. Since then many facets of homeostatic regulation has been discovered and alternative concept names have been suggested. For example, Mrosovsky [3] argued that the term *rheostasis* would be more appropriate since there is often a change in a defended set-point, for example, the elevated (and controlled) temperature when we are running a fever. He further argues (see [3], ch. 1) that homeostasis has often been equated to a single negative feedback loop. The term *allostasis* [4–6] was introduced to focus on changing environmental conditions, feedforward loops, and on the control mechanisms which deviate from a simple negative feedback loop with a single set-point [5]. With respect to circadian adaptation and anticipation mechanisms Moore-Ede [7] coined the term *predictive homeostasis*. As adaptation mechanisms are highly dynamic Lloyd [8] argued for the use of the term *homeodynamics* instead of homeostasis. While all these aspects point to important properties of homeostatic regulation, we agree with Carpenter that the term *homeostasis* still stands as an unified approach [9]. We believe, that when multiple feedback and feedforward loops are studied theoretically in more detail, many of the above mentioned homeostatic facets can be accounted for, such as rheostatic control can be observed in a model of p53 regulation upon variable stress conditions [10].

In this paper we explore the influence of background perturbations on a set of eight basic negative feedback motifs [11] with integral control. Integral control is a control-engineering concept [12], which allows a controlled variable to reset precisely at its set-point when step perturbations are applied. In biochemical systems several kinetic requirements have been identified which lead to integral control. Among them we have zero-order kinetics in the removal of the manipulated (controller) variable [11,13], antithetic control in which two controller variables are removed by second-order [14,15] or enzyme [16] kinetics, or a (first-order) autocatalytic synthesis combined with first-order removal kinetics of the manipulated variable [17–19]. When dealing with the different basic controller motifs we will introduce integral control mostly by zero-order kinetics, but also by antithetic control (see ‘Results and discussion’ below).

## Psychophysical laws

We will use the concept of a “just noticeable perturbation” (alternatively “just noticeable difference” or “threshold”) in order to compare computational results with experiments.

### Weber’s law

Ernst Heinrich Weber [20,21] found that the human perception of a just noticeable difference *dw*= *w*′−*w* between a reference weight *w* and a slightly heavier weight *w*′ is approximately proportional to the reference weight *w*, i.e.,

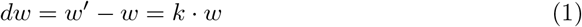

with *k* being a constant. Weber’s law implies a linear relationship between a just noticeable perturbation (threshold perturbation) and an applied background perturbation. It was Gustav Fechner [22] who made Weber’s law public and gave it its name, but expanded the perception of a just noticeable difference *dw* to *dp*=*dw/w* (termed by Fechner as *Contrast*) and stated its logarithmic form, i.e.,

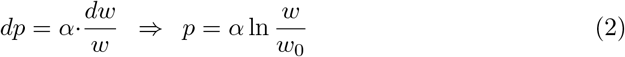

where *α* and *C* are constants and w is now a general stimulus. The logarithmic form of Eq 2 is termed as *Fechner’s law*.

### Stevens’ power law

Stevens [23] suggested (and revived) a power-law formulation between the magnitude of a sensation/perception *p* and its stimulus *s*, i.e.

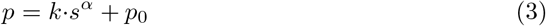

where *k, α*, and *p*_0_ are constants depending respectively on the units used and the type of stimulation. MacKay [24] suggested a model of perceived intensities by an adaptive “counterbalancing” response mechanism. This “negative feedback” approach enabled MacKay to make connections between the Weber-Fechner law and Stevens’ law. In a model of retinal light adaptation we will show that Stephens’ power law or Weber’s law are followed dependent whether the background perturbation range is either low or high, respectively.

## Materials and methods

### Calculations and parameter estimations

Computations were performed by using the Fortran subroutine LSODE [25]. Graphical results were generated with gnuplot (www.gnuplot.info). Composite figures and additional annotations were done with Adobe Illustrator (https://www.adobe.com/)

To make notations simpler, concentrations of compounds are denoted by compound names without square brackets. Time derivatives are generally indicated by the ‘dot’ notation. For the basic feedback loops m1-m8 (next section) concentrations and rate parameter values are given in arbitrary units (au), while for the light adaptation model concentrations are in *μ*M (or nM) and time scale is in seconds (s). Rate parameters are presented as *k_i_*’s (*i*=1, 2, 3,…) irrespective of their kinetic nature, i.e. whether they represent turnover numbers, Michaelis constants, or inhibition/activation constants.

For the light adaptation model some parameter values were estimated by using gnuplot’s fit function with respect to experimental literature data. Graphical experimental data were extracted with the program GraphClick (https://macdownload.informer.com/graphclick/).

To make the computations more accessible supporting information ‘S1 Programs’ contains python scripts of Fortran results.

### Feedback motifs investigated

Fig 1 shows the investigated negative feedback loops. These are eight basic motifs (m1-m8), which divide equally into a set of inflow and outflow controllers [11]. Compound *A* is the homeostatic controlled variable, while *E* is the controller variable (or manipulated variable). Red arrows indicate a step perturbation while blue arrows represent a constant background. Dashed lines represent signaling events which lead to the activation (plus signs) or inhibition (minus signs) of target reactions.

**Fig 1.**
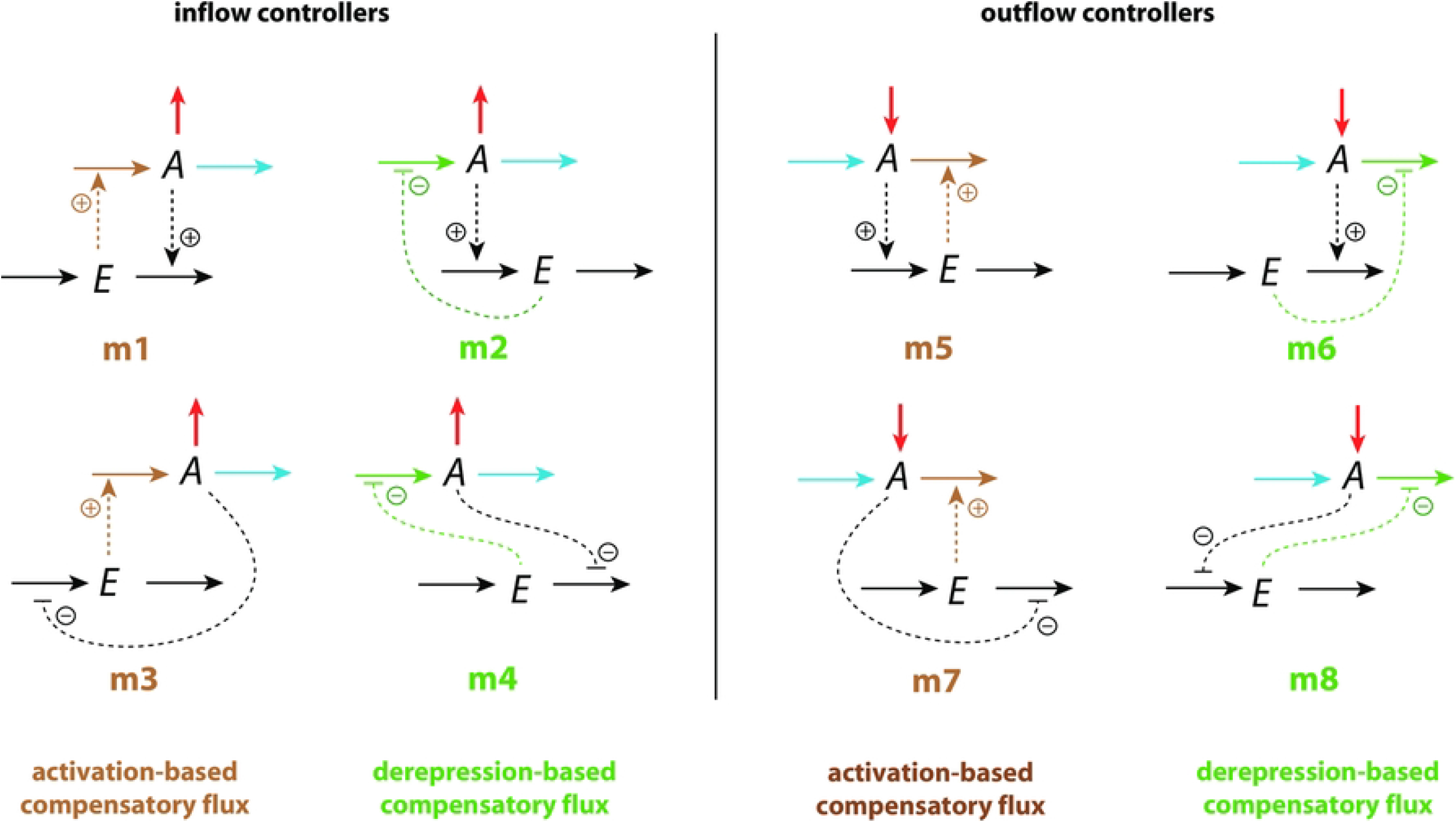
Set of basic negative feedback motifs m1-m8. Red and blue arrows indicate, respectively, a step perturbation and a constant background reaction. Integral control is implemented either by zero-order kinetics [11,13] or by antithetic control [14,16]. Outlined in brown and green we have activating or derepressing compensatory fluxes, respectively.

We have applied step perturbations, because integral controllers are generally capable to compensate them perfectly (for a proof see ch. 10.3.1 in Ref. [26]). Note however, that some feedback loop kinetics, such as in m2, are capable to oppose even rapidly increasing perturbations, such as hyperbolic growth [27,28].

In the inflow controllers m1-m4 the manipulated variable *E* leads to the increase of a compensatory inflow flux either by direct activation (brown plus signs) or by derepression (green minus signs) and thereby opposing the step perturbations which remove *A* (red arrows). In the outflow controllers m5-m8 the compensatory (outflow) flux compensates step perturbations (red arrows) which increase *A* [29].

## Results and discussion

We have analyzed the eight controller schemes (Fig 1) with regard to step perturbations at different but constant backgrounds. Fig 2 shows the two idealized responses. In panel (a) the resetting for inflow controllers is shown. In this case a step perturbation removes the controlled variable A and temporarily decreases it. Panel (b) shows the behavior of an outflow controller. When integral control is operative the controllers will defend the set-point of *A* (*A_set_*) and move the level of A during the on-going step perturbation precisely back to *A_set_*.

**Fig 2.**
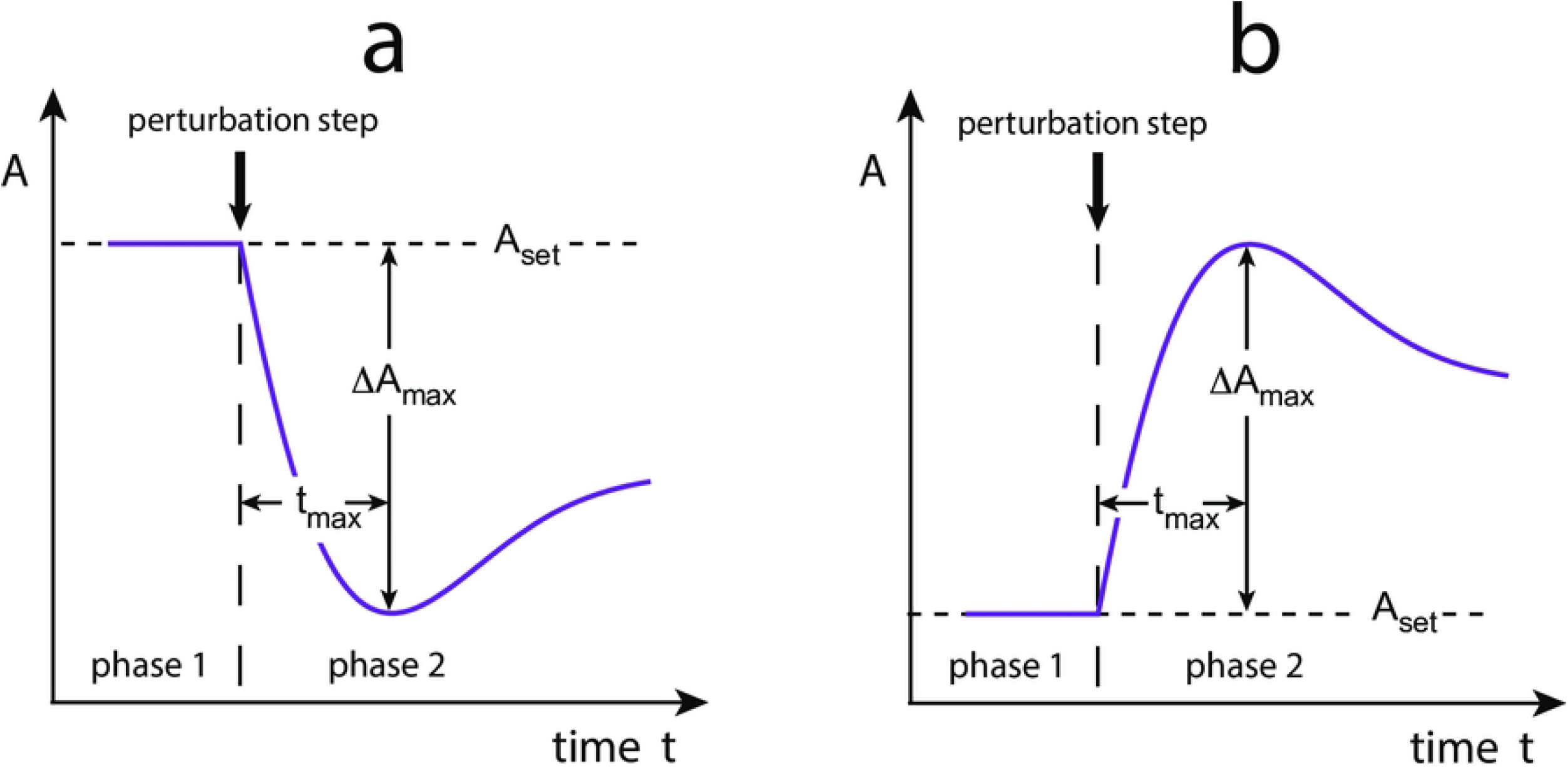
Idealized response kinetics of (a) inflow and (b) outflow controllers upon step perturbations. Indicated are the set-point of the controlled variable *A*, *A_set_*, the maximum excursion of *A*, Δ*A_max_*. *t_max_* is the time between the start of the perturbation until Δ*A_max_* is reached.

The *resetting period* is rather loosely defined as the time required to reach *A_set_* after a step perturbation has been applied. Fig 2 also indicates the parameter Δ*A_max_*, which is the maximum excursion of A after the applied step. *t_max_* is the time the controller needs to reach Δ*A_max_* after the perturbation has been applied.

We found that the controllers’ response kinetics split into two classes independent whether they are inflow or outflow controllers. In both classes an increase of a background reaction leads to a reduced excursion Δ*A_max_*. In the class where the compensatory flux is based on activation (controllers m1, m3, m5, and m7; outlined in brown in Fig 1), the controllers slow down in their resetting with increasing backgrounds and decreasing *t_max_* values. In the other class, when compensatory fluxes are based on derepression, the controllers show an accelerated resetting (controllers m2, m4, m6, and m8; outlined in green in Fig 1).

In the following we describe in more detail how the two classes of controllers differ in their resetting behavior.

### Controllers with activated compensatory fluxes

We show here the results for motifs m1 and m7. The supporting information ‘S1 Text’ shows corresponding details for controllers m3 and m5.

#### Controller m1

In the m1 controller the compensatory flux *j*_3_=*k*_3_·*E* is activated by *E* while *A* activates the removal of *E* (Fig 3). Step-wise perturbations removing *A* are mediated by *k*_2_ while *k*_4_ is a constant background outflow. For simplicity, we assume that activation kinetics are first-order with respect to the concentration of the activating species.

**Fig 3.**
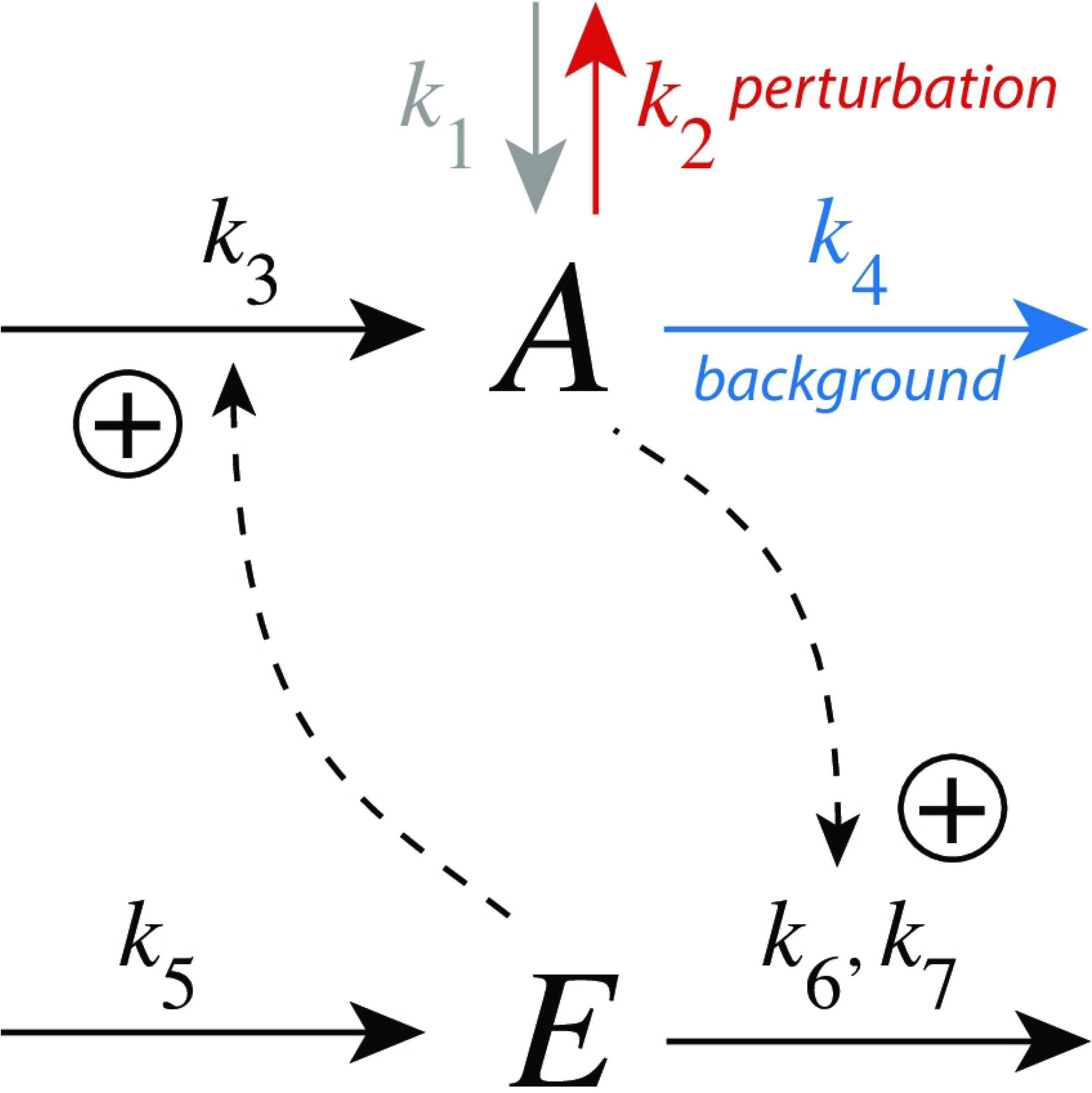
Inflow controller m1 with integral control implemented as a zero-order Michaelis-Menten (MM) type removal of *E*. *k*_2_ undergoes a step perturbation, while *k*_4_ is a constant background reaction. *k*_6_ and *k*_7_ are MM parameters analogous to *V_max_* and *K_M_*, respectively. In the calculations the grayed-out rate constant *k*_1_ will be set to zero.

The rate equations are:

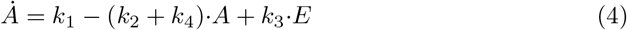

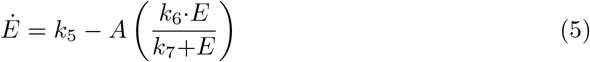

Integral control is incorporated by a zero-order kinetic removal of *E*, i.e. *E*/(*k*_7_+*E*) ≈ 1, with the result that *E* becomes proportional to the integrated error *ε* =*A_set_* – *A*:

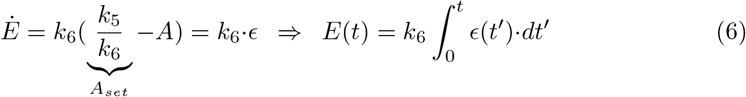

Fig 4 shows the response kinetics of the m1 controller with set-point *A_set_*=3.0. Panel (a) shows the concentration of *A* as a function of time when a *k*_2_ step 1→5 is applied. Clearly, Δ*A_max_* (see definition in Fig 2) decreases with increasing background *k*_4_. Typically for controllers where the compensatory flux is based on activation, we observe that for increased backgrounds the resetting period is lengthend. Despite the increase in the resetting period the inset in panel (a) shows that the controller is fully operational and is able to defend its set-point.

**Fig 4.**
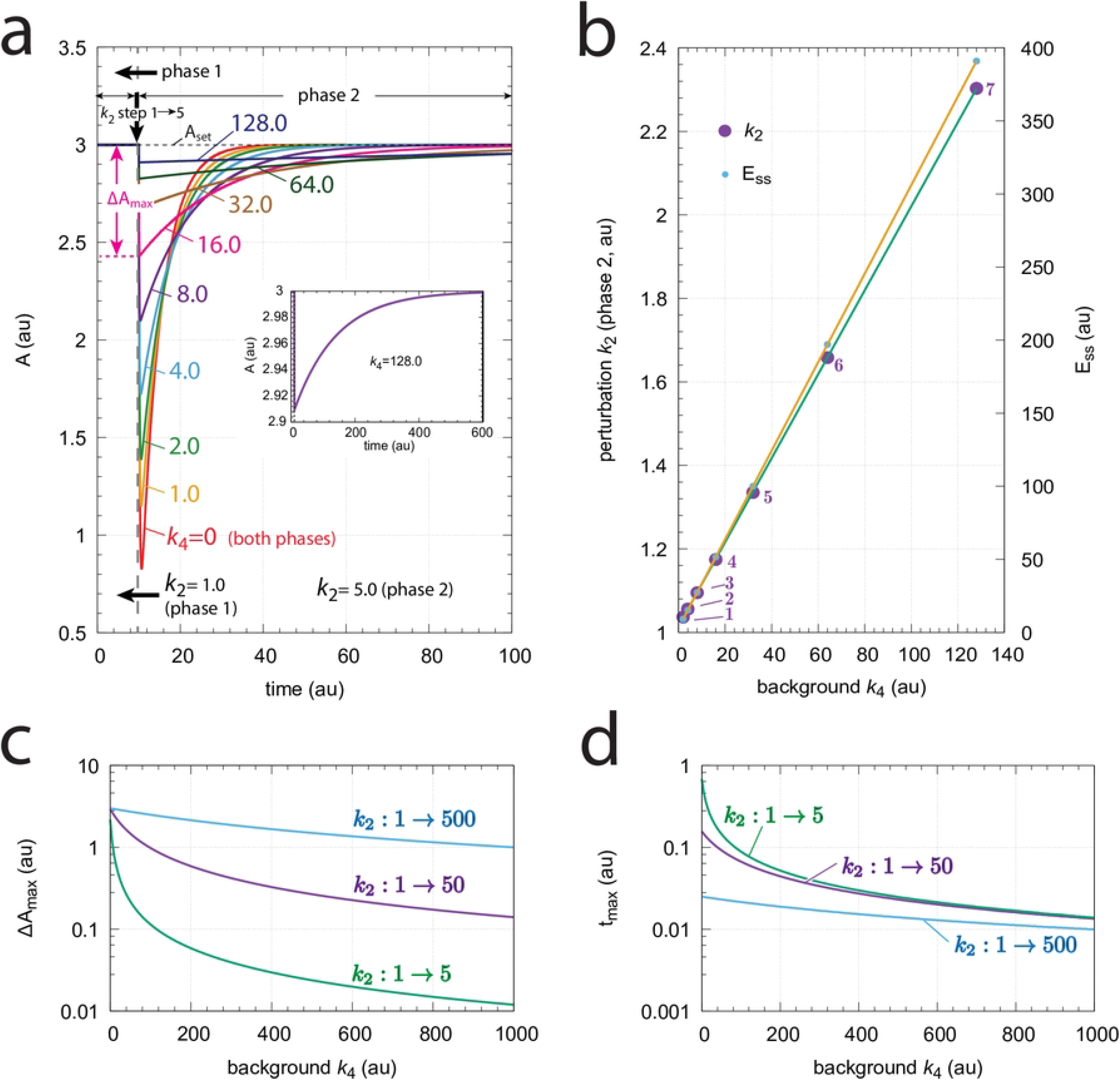
Response kinetics and relationship to Weber’s law in the m1 controller (Fig 3). The set-point of *A* is 3.0. (a) Step-wise increase of *k*_2_ from 1.0 to 5.0 at time *t*=10 at different but constant backgrounds *k*_4_ (0-128.0, phases 1 and 2). Note the successive decrease in the maximum excursion of *A* (Δ*A_max_*) with slowed-down A resetting kinetics as *k*_4_ backgrounds increase. Δ*A_max_* for *k*_4_=16.0 is indicated. The inset shows that even at high backgrounds the controller is fully operative. Rate constants (in au): *k*_1_=0.0, *k*_2_=1.0 (phase 1), *k*_2_=5.0 (phase 2), *k*_3_=1.0, *k*_4_ variable, *k*_5_=3.0, *k*_6_=1.0, *k*_7_=1×10^-6^. Initial concentrations (in au): *A*_0_=3.0, *E*_0_=3.0 (*k*_4_=0); *A*_0_=3.0, *E*_0_=6.0 (*k*_4_=1); *A*_0_=3.0, *E*_0_=9.0 (*k*_4_=2); *A*_0_=3.0, *E*_0_=15.0 (*k*_4_=4); *A*_0_=3.0, *E*_0_=27.0 (*k*_4_=8); *A*_0_=3.0, *E*_0_=51.0 (*k*_4_=16); *A*_0_=3.0, *E*_0_=99.0 (*k*_4_=32); *A*_0_=3.0, *E*_0_=195.0 (*k*_4_=64); *A*_0_=3.0, *E*_0_=387.0 (*k*_4_=128). The inset shows the full adaptation response when *k*_4_=128.0 (b) Relationship to Weber’s law: When perturbation *k*_2_ in phase 2 is adjusted such that the maximum (just noticeable) excursion in *A* is 0.03 (i.e. 1% of *A_set_*) then both *k*_2_ and the “perception” *E_ss_* are linear functions of different but constant backgrounds *k*_4_. Rate constants and initial concentrations as in (a), except that *k*_2_ in phase 2 has the following values: **1**, *k*_2_ = 1.0367 (*k*_4_ = 2); **2**, *k*_2_ = 1.0559 (*k*_4_ = 4); **3**, *k*_2_ = 1.0950 (*k*_4_ = 8); **4**, *k*_2_ = 1.1745 (*k*_4_ = 16); **5**, *k*_2_ = 1.3350 (*k*_4_ = 32); **6**, *k*_2_ = 1.6581 (*k*_4_ = 64); **7**, *k*_2_ = 2.3030 (*k*_4_ = 128). (c) Δ*A_max_* as a function of background *k*_4_ at three different *k*_2_ steps. (d) *t_max_* as a function of background *k*_4_ at three different *k*_2_ steps. Rate constants are as in panel (a), except for *k*_2_ and *k*_4_. Initial concentrations are the steady state values of *A* and *E* prior to the step in *k*_2_.

Fig 4b shows the response kinetics related to Weber’s law when probing a “just noticeable” excursion in Δ*A* of 0.03 (1% of *A_set_* =3.0) by applying appropriate *k*_2_ values in phase 2. We observe that the different *k*_2_ values (in phase 2) and the corresponding steady-state values of *E* (*E_ss_*) are linear functions of the background *k*_4_.

Figs 4c and d show the values of Δ*A_max_* and *t_max_* for three different *k*_2_ steps with increasing backgrounds *k*_4_. Reflecting the behavior from Fig 4a panel (c) shows that Δ*A_max_* values decrease monotonically as background increases, but that the magnitude of Δ*A_max_* depends on the size of the applied step. Despite that the resetting period increases with increasing backgrounds we observe that *t_max_* decreases with increasing *k*_4_ (panel (d)). The increase of the resetting period at increased *k*_4_ levels can be explained by the high steady state levels of *E* in phase 1 when *k*_4_ backgrounds become high and that the system needs more time to reach the steady state of *E* in phase 2 by zero-order kinetics.

#### Controller m7

m7 is an outflow controller which opposes inflow perturbations *k*_1_ at different background reactions *k*_3_ by *E*-activation of the compensatory flux *j*_4_ (=*k*_4_·*A*·*E*). The negative feedback is closed by inhibiting the removal of *E* through *A* (Fig 5). The rate equations are

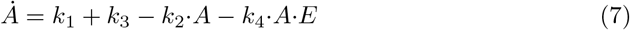

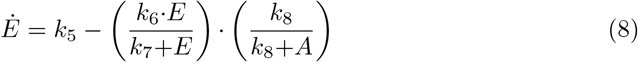

**Fig 5.**
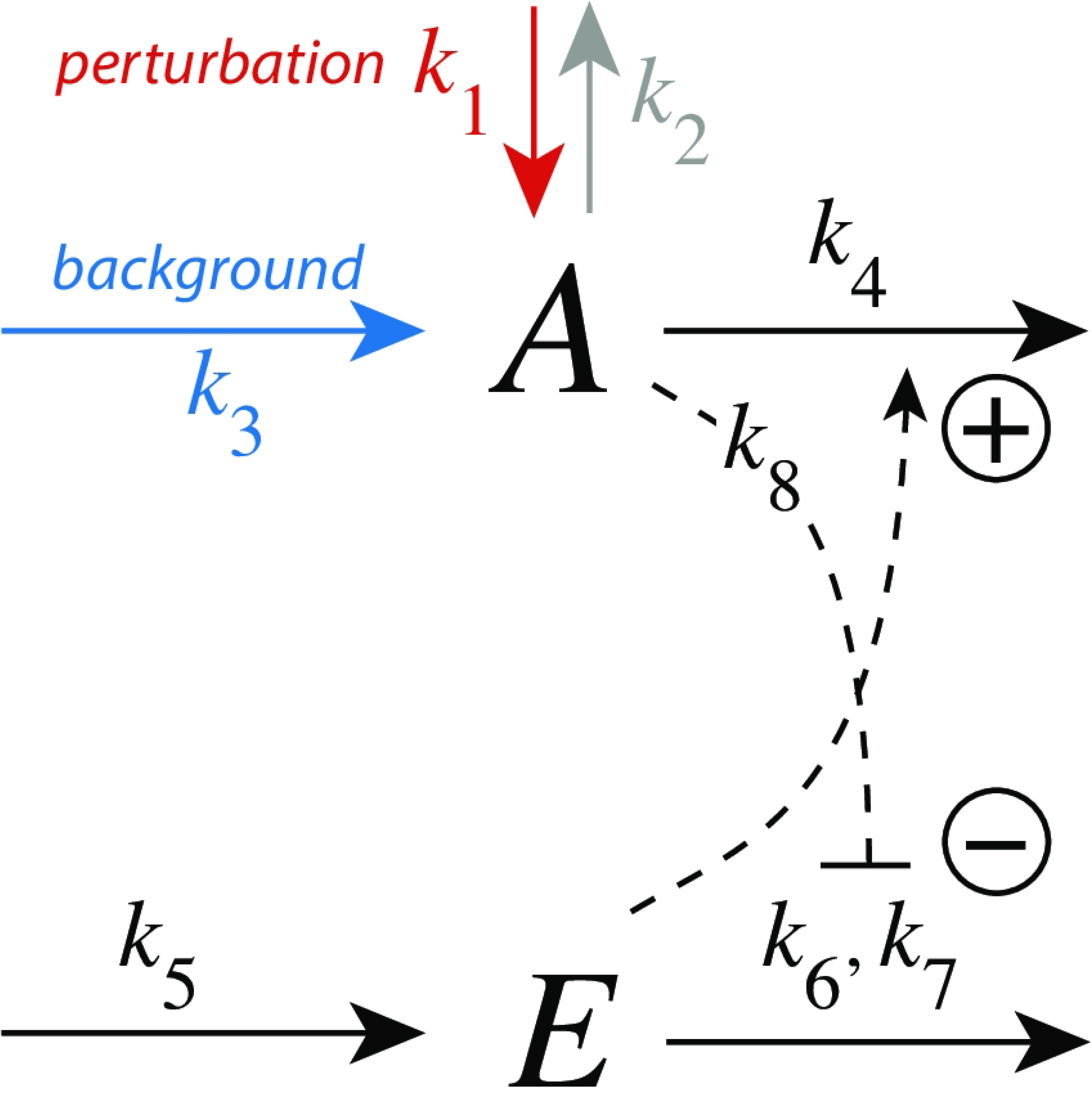
Outflow controller motif m7 with integral control implemented as a zero-order Michaelis-Menten (MM) type degradation of *E*. The perturbation *k*_1_ changes step-wise (1.0→5.0), while *k*_3_ is a constant background. Rate constant *k*_8_ is an inhibition constant. *k*_6_ and *k*_7_ are MM parameters analogous to *V_max_* and *K_M_*, respectively. In the calculations the grayed-out rate constant *k*_2_ is, for the sake of simplicity, set to zero.

The set-point for *A* is calculated from the steady-state condition of Eq 8 by using zero-order degradation of *E*, i.e. *E*/(*k*_7_+*E*)≈1.

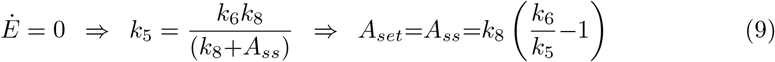

Fig 6 shows the response kinetics of the m7 controller. Since the controller opposes inflow perturbations excursions of *A* are above the set-point *A_set_* (=3.0). Panel a shows the slowed-down responses during the resetting in phase 2 as background *k*_3_ increases. The inset shows that the controller is still operative even at the highest *k*_3_ and slowest resetting. Panel b shows that a *k*_1_ step perturbation which results in a just noticeable maximum excursion Δ*A_max_* of 0.03 (1% of *A_set_*) increases, together with the corresponding steady state *E_ss_* values in phase 2, linearly with the background *k*_3_. Δ*A_max_* in creases with increasing *k*_1_ step (Fig 6c), while for a given background we find, somewhat surprisingly, that *t_max_* is independent on the magnitude of the *k*_1_ step (Fig 6d). Both Δ*A_max_* and *t_max_* decrease monotonically with increasing background *k*_3_.

**Fig 6.**
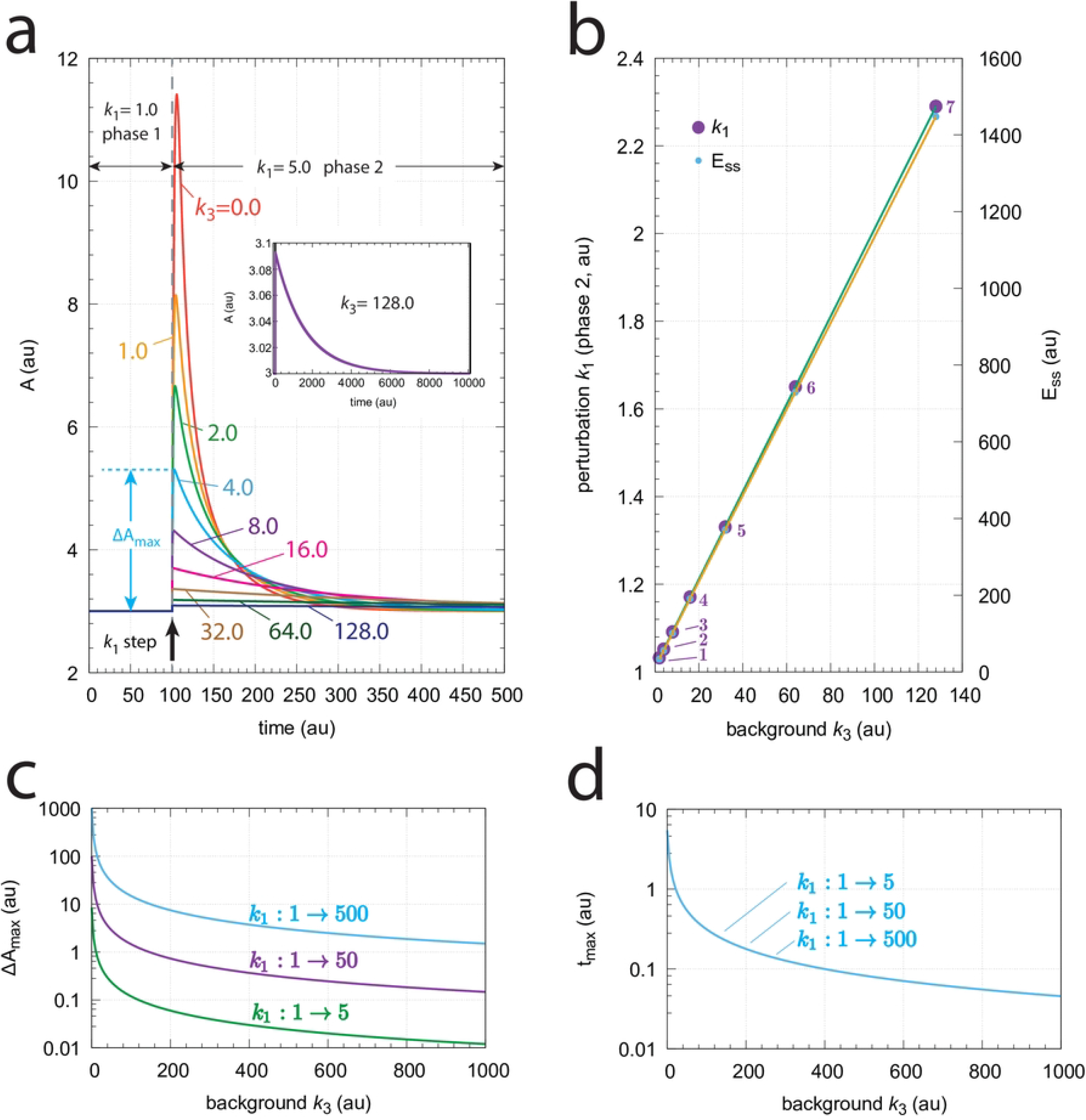
Response kinetics and relationship to Weber’s law in the m7 controller (Fig 5). The set-point of *A* is 3.0. (a) Step-wise increase of *k*_1_ from 1.0 to 5.0 at time *t*=100 at different and constant background perturbations *k*_3_ (0-128.0, applied in phases 1 and 2). Note the successive decrease in the maximum excursion of *A* (Δ*A_max_*) with slowed-down A resetting kinetics as *k*_3_ values increase. Δ*A_max_* for *k*_3_=4.0 is indicated. Rate constants (in au): *k*_1_=1.0, *k*_2_=0.0 (phases 1 and 2), *k*_1_=5.0 (phase 2), *k*_3_ variable, *k*_4_=0.03, *k*_5_=1.0, *k*_6_=31.0, *k*_7_=1×10^-6^, *k*_8_=0.1. Initial concentrations (in au): *A*_0_=3.0, *E*_0_=11.11 (*k*_3_=0); *A*_0_=3.0, *E*_0_=22.22 (*k*_3_=1); *A*_0_=3.0, *E*_0_=33.33 (*k*_3_=2); *A*_0_=3.0, *E*_0_=55.55 (*k*_3_=4); *A*_0_=3.0, *E*_0_=100.0 (*k*_3_=8); *A*_0_=3.0, *E*_0_=188.89 (*k*_3_=16); *A*_0_=3.0, *E*_0_=366.67 (*k*_3_=32); *A*_0_=3.0, *E*_0_=722.22 (*k*_3_=64); *A*_0_=3.0, *E*_0_=1433.33 (*k*_3_=128). The inset shows the full adaptation response when *k*_3_=128.0 (b) Relationship to Weber’s law: When perturbation *k*_1_ in phase 2 is adjusted such that the maximum (just noticeable) excursion Δ*A_max_* is 0.03 (i.e. 1% of *A_set_*) then both *k*_1_ and the “perception” *E_ss_* are linear functions of the background *k*_3_. Rate constants and initial concentrations as in (a), except that *k*_1_ in phase 2 has the following values: **1**, *k*_1_ = 1.0325 (*k*_3_ = 2); **2**, *k*_1_ = 1.0520 (*k*_3_ = 4); **3**, *k*_1_ = 1.0914 (*k*_3_ = 8); **4**, *k*_1_ = 1.1709 (*k*_3_ = 16); **5**, *k*_1_ = 1.3306 (*k*_3_ = 32); **6**, *k*_1_ = 1.6503 (*k*_3_ = 64); **7**, *k*_1_ = 2.2900 (*k*_3_ = 128). (c) Δ*A_max_* values as a function of background *k*_3_ for three step perturbations in *k*_1_. Note that the three curves are congruent, i.e., their identical shape can be precisely moved onto each other. (d) *t_max_* as a function of background *k*_3_. For a given background *t_max_* is practically the same and independent of the three *k*_1_ steps.

We explain the delay in the resetting of *A* for large *k*_3_ backgrounds as the increased time needed to change the high steady state values of *E* from phase 1 to its new steady state in phase 2 after the step.

### Controllers with compensatory fluxes based on derepression

We show here the results for controllers m2 and m8. Corresponding results for m4 and m6 are given in supporting information ‘S2 Text’.

#### Controller m2

In the m2 controller scheme (Fig 7) activation of *E* by *A* is proportional to the concentration of *A*, while the inhibition term on the compensatory flux is formulated as *k*_8_/(*k*_8_+*E*). The rate equations are:

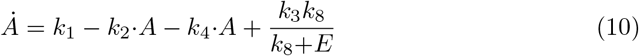

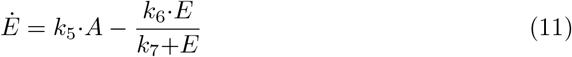

**Fig 7.**
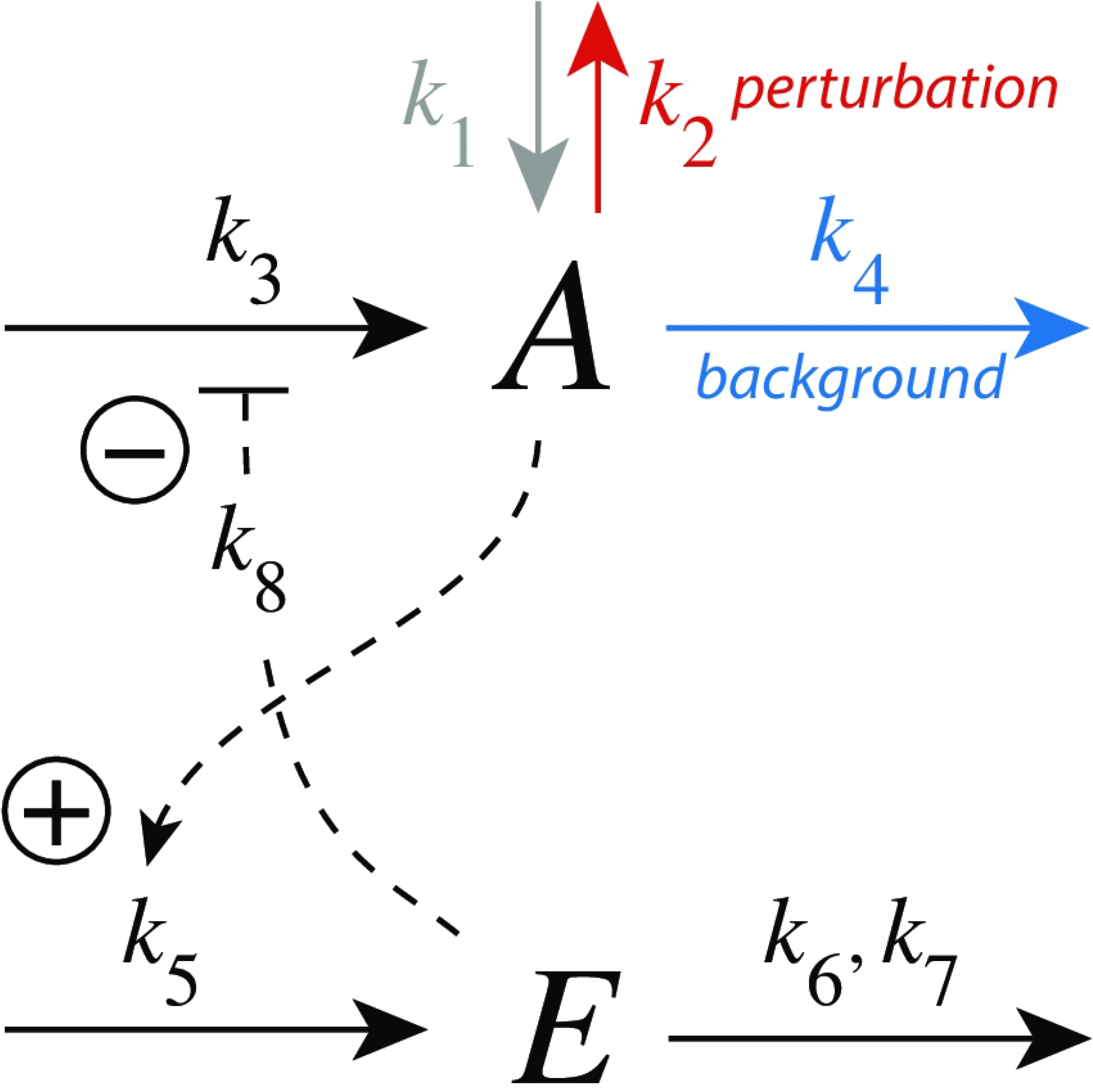
Controller motif m2 with integral control implemented as a zero-order Michaelis-Menten (MM) type degradation of E. Rate constant *k*_2_ undergoes a step-wise change (perturbation), while *k*_4_ is a (constant) background reaction. Rate constant *k*_8_ is an inhibition constant. *k*_6_ and *k*_7_ are MM parameters analogous to *V_max_* and *K_M_*, respectively. The grayed-out rate constant *k*_1_ is set in the calculations to zero.

To achieve homeostasis in *A* a perturbation (removal) of *A* is counteracted by a decrease of *E* (“derepression”), which increases the compensatory flux *j*_3_=*k*_3_*k*_8_/(*k*_8_+*E*) and moves, in the presence of integral control, A to its set-point.

The set-point of *A* (*A_set_*) is determined how integral control is implemented in the feedback loop. In Fig 7 we use zero-order kinetics with respect to the removal of *E*, i.e. *k*_7_ ≪ *E_ss_*. This implies that the steady state of *A* is also the set-point of *A* (*A_set_*) and is given as the ratio *k*_6_/*k*_5_, i.e.

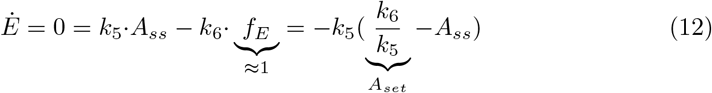

with *f_E_* = *E*/(*k*_7_+*E*)≈1.

Fig 8a shows the response for step-wise changes in *k*_2_ from 1.0 (phase 1) to 5.0 (phase 2) at different but constant background perturbations *k*_4_. Typically for derepression controllers is both the decrease of Δ*A_max_* at increasing backgrounds when a constant step perturbation is applied and a *decreasing* response time.

**Fig 8.**
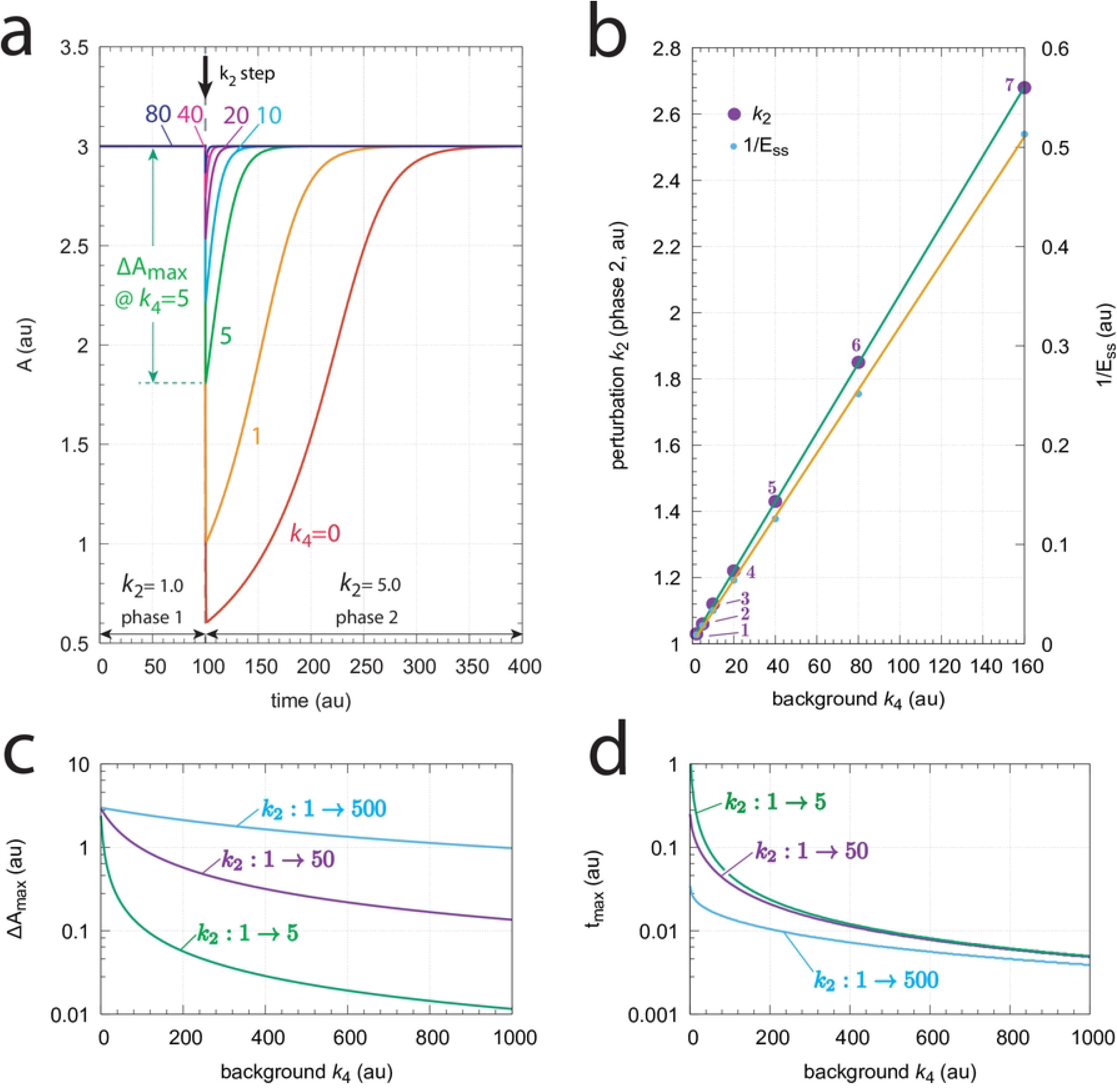
Response kinetics and relationship to Weber’s law in the m2 controller (Fig 7). The set-point of *A* is *A_set_*=3.0 (a) Step-wise increase of *k*_2_ from 1.0 to 5.0 at time *t*=100 at different and constant background perturbations *k*_4_ (0-80). The maximum excusion in *A*, Δ*A_max_*, for *k*_4_=5 is indicated. Note the successive decrease in Δ*A_max_* and the more rapid resetting of *A* at increased *k*_4_ values. Rate constants (in au): *k*_1_=0.0, *k*_2_=1.0 (phase 1), *k*_2_=5.0 (phase 2), *k*_3_=1×10^4^, *k*_4_ variable, *k*_5_=1.0, *k*_6_=3.0, *k*_7_=1×10^-6^, *k*_8_=0.1. Initial concentrations (in au): *A*_0_=3.0, *E*_0_=333.23 (*k*_4_=0); *A*_0_=3.0, *E*_0_=166.62 (*k*_4_=1); *A*_0_=3.0, *E*_0_=55.46 (*k*_4_=5); *A*_0_=3.0, *E*_0_=30.20 (*k*_4_=10); *A*_0_=3.0, *E*_0_=15.77 (*k*_4_=20); *A*_0_=3.0, *E*_0_=8.03 (*k*_4_=40); *A*_0_=3.0, *E*_0_=4.02 (*k*_4_=80). (b) Relationship to Weber’s law: in phase 2 the perturbation *k*_2_ and (1/*E_ss_*) are linear functions of the background perturbation *k*_4_ when the “just noticable difference” Δ*A_max_* is 0.03. Rate constants and initial concentrations as in (a), except that *k*_2_ in phase 2 has the following values: **1**, *k*_2_ = 1.0314 (*k*_4_ = 2); **2**, *k*_2_ = 1.0627 (*k*_4_ = 5); **3**, *k*_2_ = 1.1150 (*k*_4_ = 10); **4**, *k*_2_ = 1.2195 (*k*_4_ = 20); **5**, *k*_2_ = 1.4285 (*k*_4_ = 40); **6**, *k*_2_ = 1.8465 (*k*_4_ = 80); **7**, *k*_2_ = 2.6820 (*k*_4_ = 160). (c) Monotonic decrease of Δ*A_max_* as a function of background *k*_4_ for three different steps. At constant background Δ*A_max_* increases with increasing step size. (d) t_max_ decreases monotonically with increasing backgrounds *k*_4_. At constant background t_max_ decreases with increasing step size. Rate constants in panels (c) and (d) are the same as for panel (a), apart from *k*_2_ and *k*_4_. Initial concentrations were taken as the steady state values of *A* and *E* at the different backgrounds *k*_4_ prior to the applied step in *k*_2_.

We were interested to see how the m2 controller would respond when a just noticeable excursion in *A* (Δ*A_max_*) was applied for different background perturbations *k*_4_. For that purpose we determined in phase 2 the steady state values of *E* and the *k*_2_ values when the excursion of *A* was 1% of *A_set_* (=3.0), i.e. Δ*A_max_*=0.03. Fig 8b shows that (1/*E_ss_*) and *k*_2_ increase linearly with increasing *k*_4_, a manifestation of Weber’s law. In this view, (1/*E_ss_*) could be interpreted as a “perceived” variable.

#### Controller m2 with antithetic integral control

Since we will later include a bimolecular (antithetic) mechanism in retinal photoadaptation, which shows robust perfect adaptation, we include here scheme m2 with antithetic [14,19] control (Fig 9).

The rate equations are:

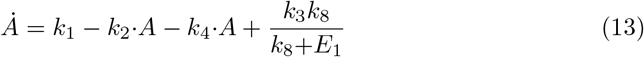

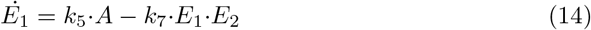

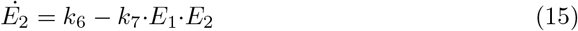

**Fig 9.**
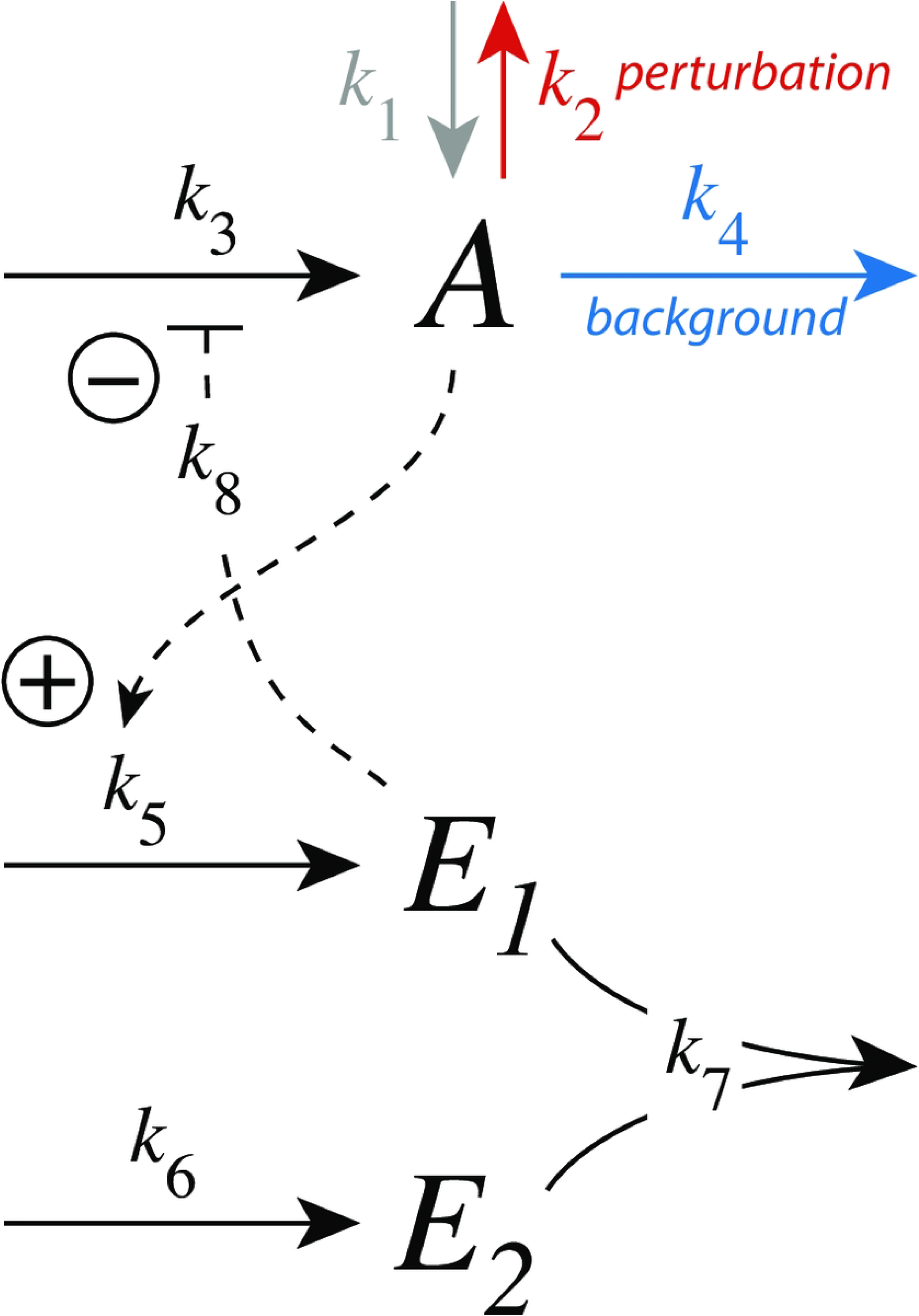
Controller motif m2 with antithetic integral control. Here, antithetic control is implemented as a bimolecular second-order reaction which removes the two controller molecules *E*_1_ and *E*_2_. See text on how *A*’s set-point is calculated.

From the steady-state conditions for *E*_1_ (*k*_5_·*A_ss_*=*k*_7_·*E*_1_·*E*_2_) and *E*_2_ (*k*_6_=*k*_7_·*E*_1_·*E*_2_) the set-point for *A* (*A_set_*) is given by:

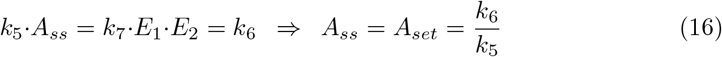

In many respects robust perfect adaptation by zero-order or bimolecular (antithetic) kinetics, i.e., *E* (Eq 12) and *E*_1_ (Eq 14) behave dynamically identical. In fact, both *E* and *E*_1_ show zero-order kinetics with respect to *E* and *E*_1_, respectively. In the supporting information ‘S3 Text’ we show the identical antithetic behavior of the m2 scheme when using step perturbations at various backgrounds in comparison with the above m2 calculations using zero-order kinetics.

#### Controller m8

Fig 10 shows the scheme of controller m8. The compensatory outflow flux *j*_4_=*k*_4_·*k*_9_·*A*/((*k*_9_+*E*)) and the signaling from A to E are based on derepression.

**Fig 10.**
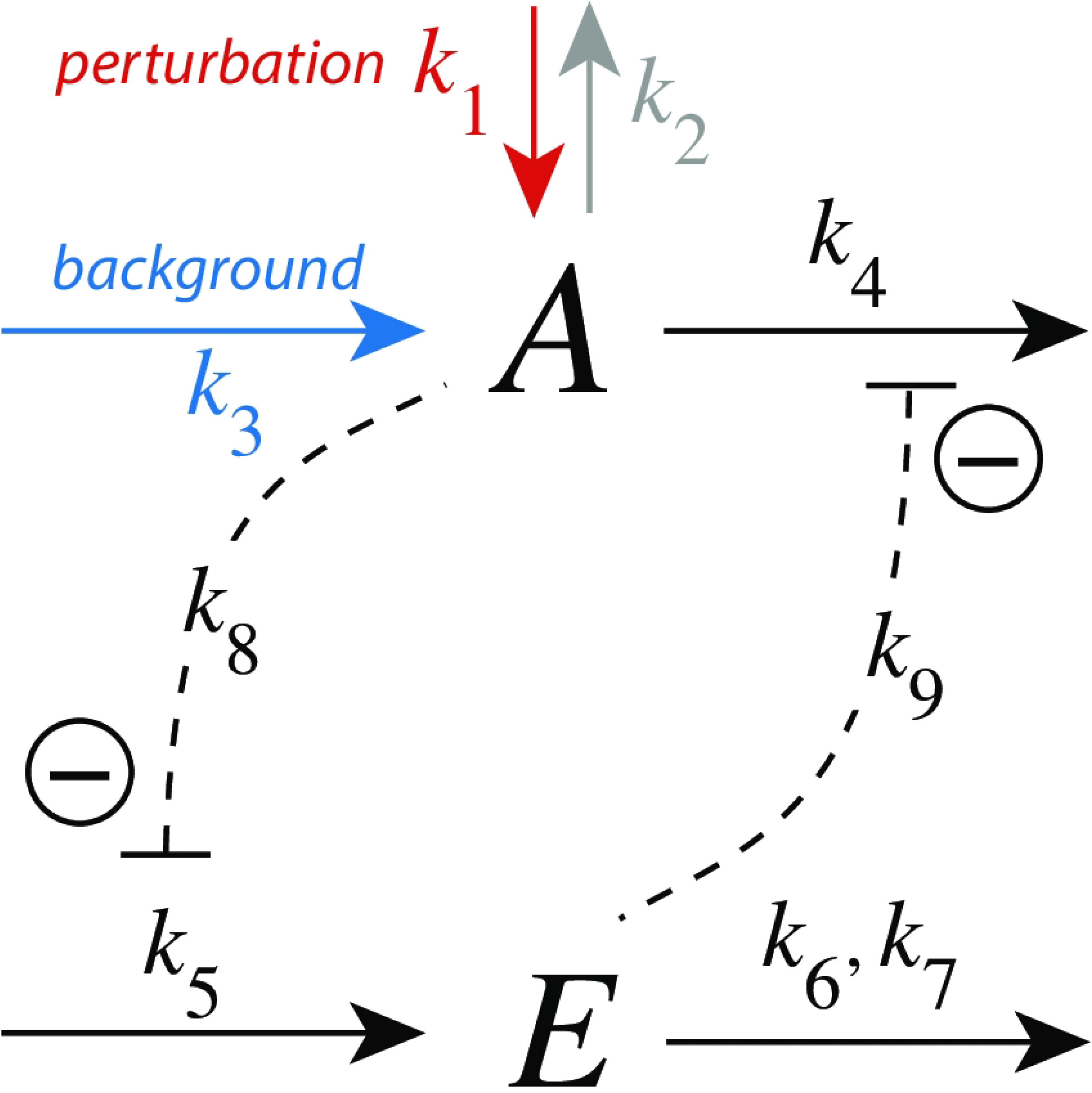
Outflow controller motif m8 with integral control implemented as a zero-order Michaelis-Menten (MM) type degradation of E. Rate constant *k*_1_ undergoes a perturbation, while *k*_3_ is a background inflow rate. *k*_8_ and *k*_9_ are inhibition constants. *k*_6_ and *k*_7_ are MM parameters analogous to *V_max_* and *K_M_*, respectively. For simplicity, the grayed-out rate constant *k*_2_ is set to zero.

The rate equations are:

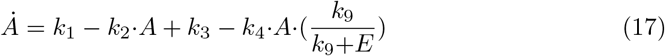

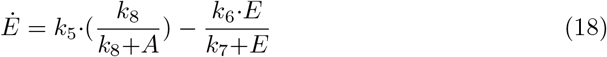

The set-point of *A* is derived from the steady-state condition *Ė*=0 together with the assumption that *E* is removed by zero-order kinetics, i.e. *E*/(*k*_7_+*E*)≈1:

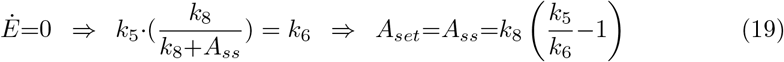

Fig 11a shows the response of the m8 derepression controller at different but constant backgrounds *k*_3_. Note the typical, more rapid, resetting when backgrounds are increased. Panel b shows the relationship to Weber’s law. When setting a “just noticeable difference” of Δ*A_max_* to 1% of the set-point of *A* (*A_set_*=3.0) the required perturbations *k*_1_ in phase 2 needed to achieve Δ*A_max_*=0.03 become a linear function of the background *k*_3_. Similarly, plotting (1/*E_ss_*) against the background is likewise linear, suggesting that (1/*E_ss_*) may be interpreted as the ‘perception” of Δ*A_max_*.

**Fig 11.**
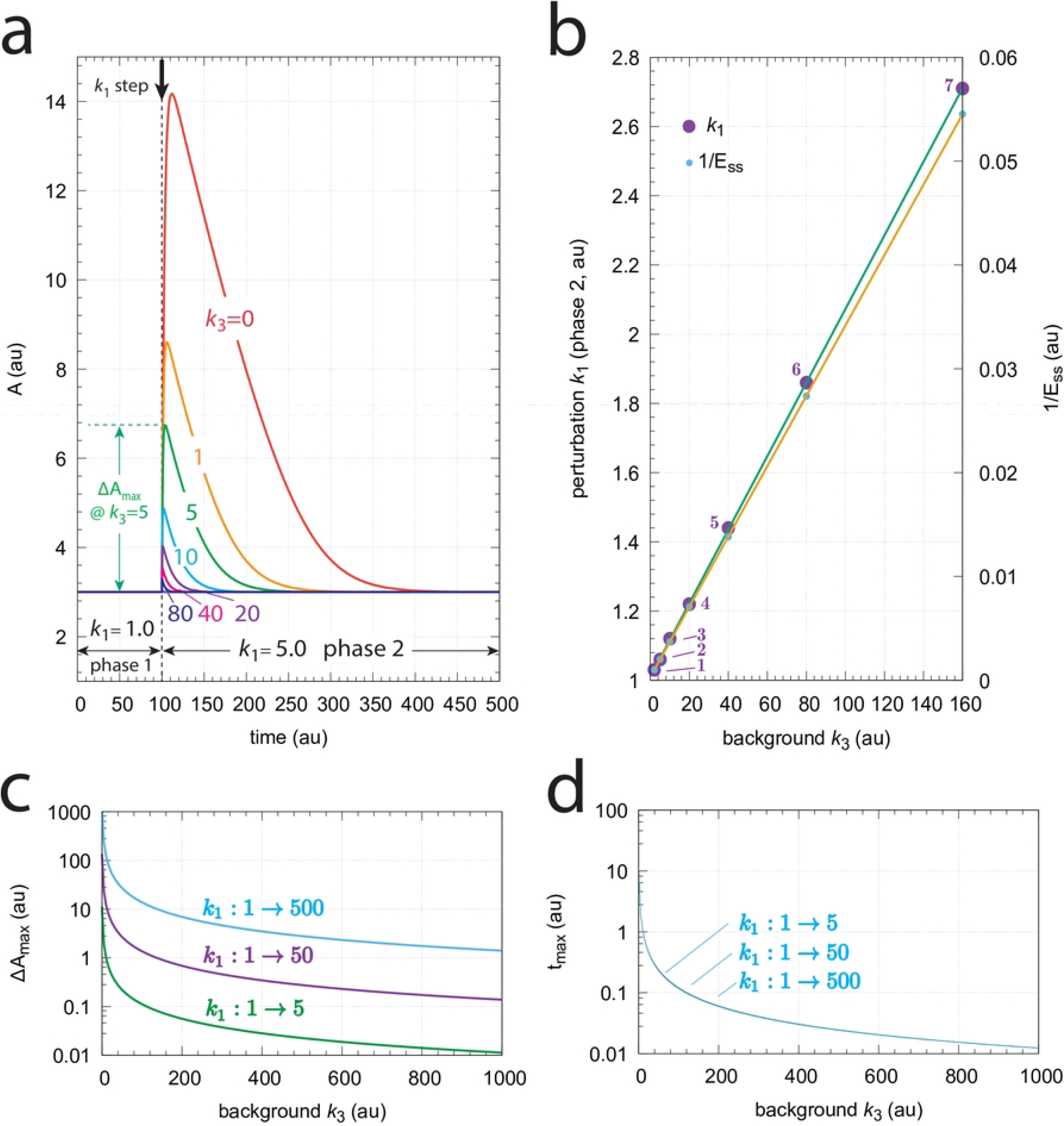
Response kinetics and relationship to Weber’s law in the m8 controller (Fig 10). The set-point of *A* is *A_set_*=3.0. (a) Step-wise increase of *k*_1_ from 1.0 to 5.0 at time *t*=100 at different and constant background perturbations *k*_3_ (0-80). Note the successive decrease in the excursion of *A* (Δ*A_max_*) and the more rapid *A* resetting to the set-point at increased *k*_3_ values. Rate constants (in au): *k*_1_=1.0 (phase 1), *k*_1_=5.0 (phase 2), *k*_2_=0.0, *k*_3_ variable, *k*_4_=1×10^4^, *k*_5_=620.0, *k*_6_=20.0, *k*_7_=1×10^-6^, *k*_8_=*k*_9_=0.1. Initial concentrations (in au): *A*_0_=3.0, *E*_0_=2999.90 (*k*_3_=0); *A*_0_=3.0, *E*_0_=1499.90 (*k*_3_=1); *A*_0_=3.0, *E*_0_=499.90 (*k*_3_=5); *A*_0_=3.0, *E*_0_=272.63 (*k*_3_=10); *A*_0_=3.0, *E*_0_=142.76 (*k*_3_=20); *A*_0_=3.0, *E*_0_=73.07 (*k*_3_=40); *A*_0_=3.0, *E*_0_=36.94 (*k*_3_=80). (b) Relationship to Weber’s law: the perturbation *k*_1_ and (1/*E_ss_*) in phase 2 are linear functions of the background perturbation *k*_3_ when *k*_1_ is adjusted such that a “just noticable difference” of Δ*A_max_*=0.03 is observed. Rate constants and initial concentrations as in (a), except that *k*_1_ in phase 2 has the following values: **1**, *k*_1_ = 1.0319 (*k*_3_ = 2); **2**, *k*_1_ = 1.0637 (*k*_3_ = 5); **3**, *k*_1_ = 1.1169 (*k*_3_ = 10); **4**, *k*_1_ = 1.2231 (*k*_3_ = 20); **5**, *k*_1_ = 1.4356 (*k*_3_ = 40); **6**, *k*_1_ = 1.8604 (*k*_3_ = 80); **7**, *k*_1_ = 2.7111 (*k*_3_ = 160). (c) Δ*A_max_* values as a function of background *k*_3_ for three step perturbations in *k*_1_. Like for the m7 controller the three curves are congruent and their shape can be moved onto each other. (d) *t_max_* as a function of background *k*_3_. For a given background *t_max_* values are practically the same independent of the three steps. Rate constants in panels (c) and (d) are the same as for panel (a), apart from *k*_1_ and *k*_3_. Initial concentrations are taken as the steady state values for *A* and *E* at the different backgrounds *k*_3_ prior to the applied step in *k*_1_.

### Model of photoreceptor adaptation

As a biological example, we found a striking analogy between the resetting kinetics of the derepression controllers m2, m4, m6, and m8 and the responses in vertebrate photoreceptors. Fig 12 shows voltage responses of a rod cell to 10 ms light flashes at different background light intensities [30]. The experiments show that increased backgrounds lead to diminished response excursions, while the resetting to the initial steady state levels were found to be faster. This behavior, a decreased sensitivity but accelerated response kinetics at increased background light intensities is considered typical for the light adaptation in vertebrate rod or cone cells [31]. When corresponding photocurrents are studied as a function of different background light levels the observed resetting behavior is close to that found for m8 or m6 controllers (for experimental data see Fig 1 in Ref [32]).

**Fig 12.**
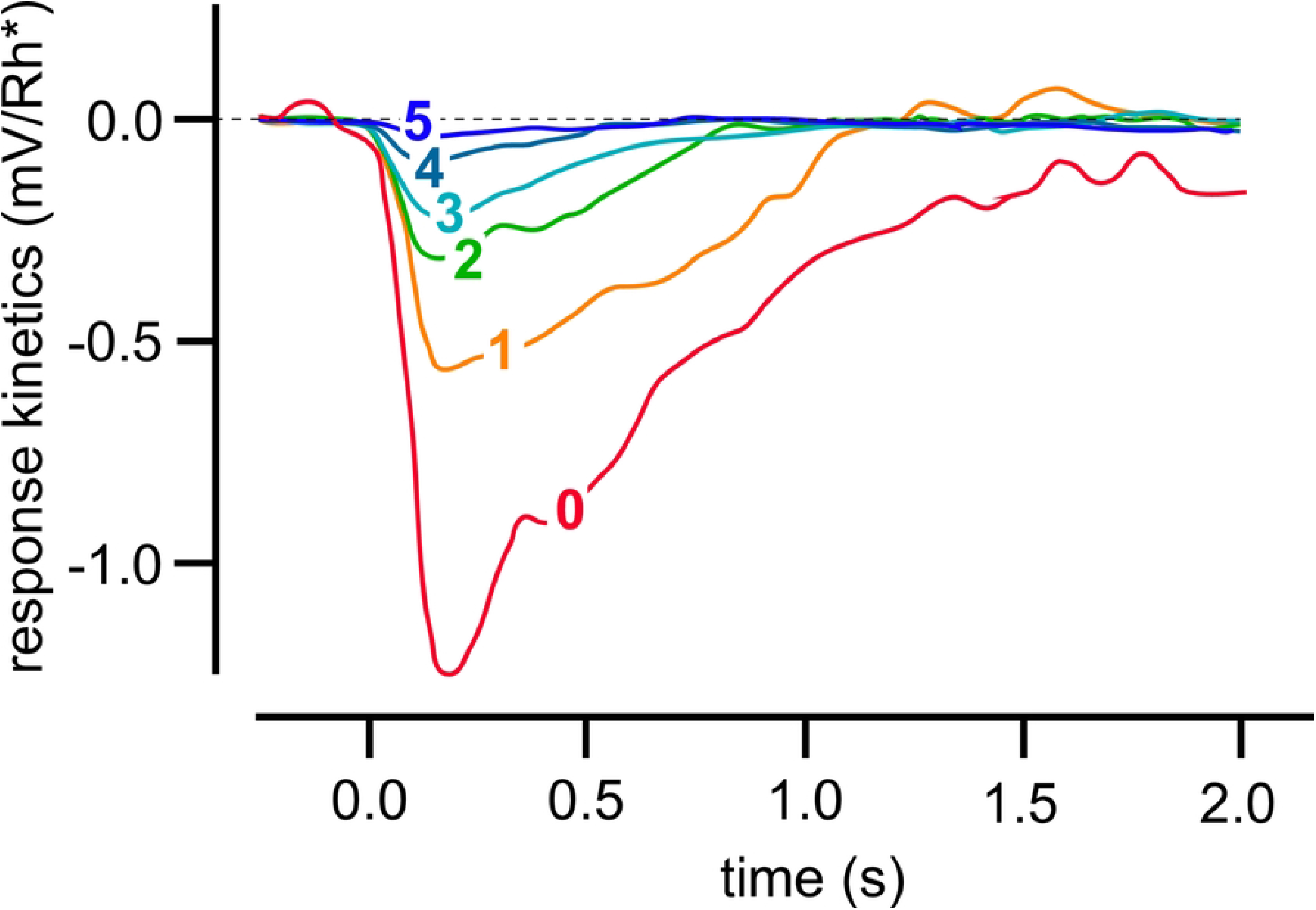
Light adaptation in a Macaque monkey’s rod cell. 10 ms light flashes were applied to different light background intensities. Background intensities (in photons *μ*m^-2^s^-1^) were: **0**, 0; **1**, 3.1; **2**, 12; **3**, 41; **4**, 84; **5**, 162. Redrawn after Fig 2A from Ref [30].

In photoreceptor cells cytosolic calcium has been found to be the major regulator in vertebrate light adaptation [33]. There, calcium takes part in a derepressing feedback loop analogous as *E* in m2. Fig 13 shows a model with its main regulatory elements. In comparison with extracellular Ca^2+^ concentrations, which are in the 10-100 mM range, cytosolic (internal) Ca^2+^ levels 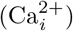 are considerably lower, around in the 100 nM range since too high cytosolic 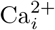 concentrations are toxic and may lead to apoptosis. Thus, despite that Ca^2+^ is a versatile cellular signal 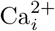 levels are tightly regulated [34]. In photoreceptor cells dark 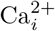 levels are in the range around 300-500 nM [33], which is sufficient to regulate photo-transduction, but at the same time low enough to avoid cytotoxic Ca^2+^ effects.

**Fig 13.**
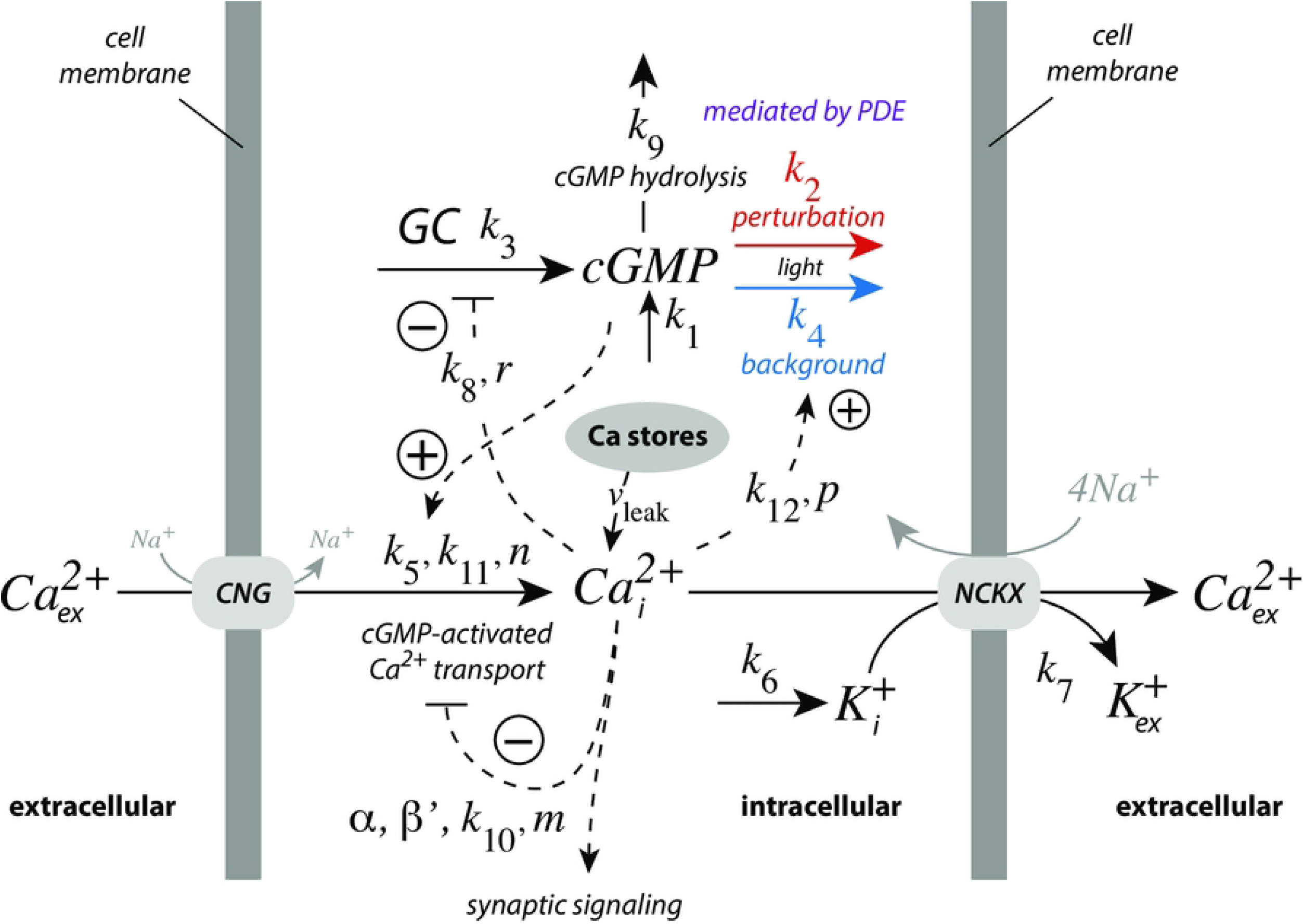
Model with the main regulatory elements of photoreceptor adaptation. Light leads to the removal of cyclic guanosine monophosphate (cGMP) by phospho-diesterases (PDE), via transducin and the activation of PDE by internal 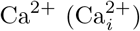. cGMP is formed by guanylate cyclase (GC). cGMP’s constitutive non-light induced hydrolysis is described by a first-order reaction with rate constant *k*_9_. GC is inhibited/derepressed by 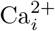. cGMP activates cyclic nucleotide-gated (CNG) channels, which leads to the inflow of Ca^2+^ into the cell, while high 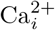 levels inhibit CNG channels. Calcium is removed from the cell by potassium-dependent sodium-calcium exchangers (NCKX). Rate equations and used rate parameter values are described in the main text. Grayed reaction arrows indicate reactions which are not included in the model.

In photoreceptor cells 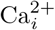 is part of a negative feedback regulation of cyclic guanosine monophosphate (cGMP), where cGMP activates the inflow of Ca^2+^ into the cytosol by cyclic nucleotide-gated (CNG) channels. Analogous to a m2 controller, 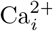 on its side inhibits guanylate cyclase (GC), which synthesizes cGMP. In addition, Ca^2+^ inhibits its inflow by CNG channels and takes part, analogous to a m5 controller, in the light-dependent removal of cGMP by activating phospho-diesterases (PDE). Potassium-dependent sodium-calcium exchangers (NCKX) pump 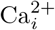 out of the cell. In the model the removal of 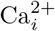 by NCKX is formulated, for the sake of simplicity, as a bimolecular second-order reaction, where K^+^ is removed together with 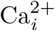, while keeping NCKX constant. For certain feedback combinations the bimolecular (or a zero-order) removal of 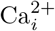 and K^+^ by NCKX will lead to robust perfect adaptation of cGMP, which is discussed below. *k*_1_ represents an inflow perturbation with respect to cGMP. We have mostly ignored *k*_1_, except in section “Roles of the feedback loops”, where *k*_1_ is used to test the homeostatic behaviors of the individual feedback loops.

The rate equations of the model are:

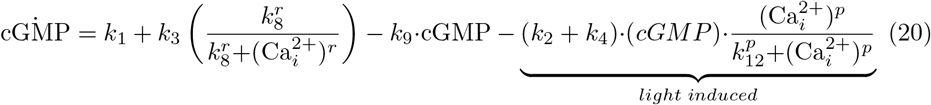

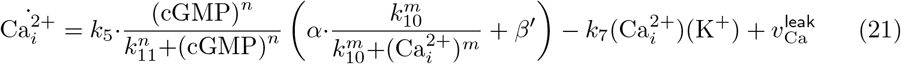

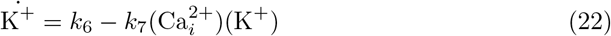

#### Estimation of model parameters

Fig 14 gives an overview of the experimental data used to estimate some of the model parameters. Panel a shows the results by Koutalos et al. (Fig 3 in [35]; see also Fig 3 in [36]), who studied the influence of Ca^2+^ on the light-stimulated PDE activity in salamander rods. The experimental data were described by the function

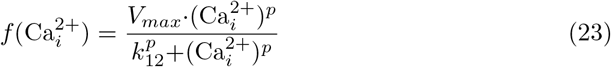

with *V_max_*=(100.01±2.53)%, *p*=0.894±0.0534, and *k*_12_=(622.612±55.01)nM.

**Fig 14.**
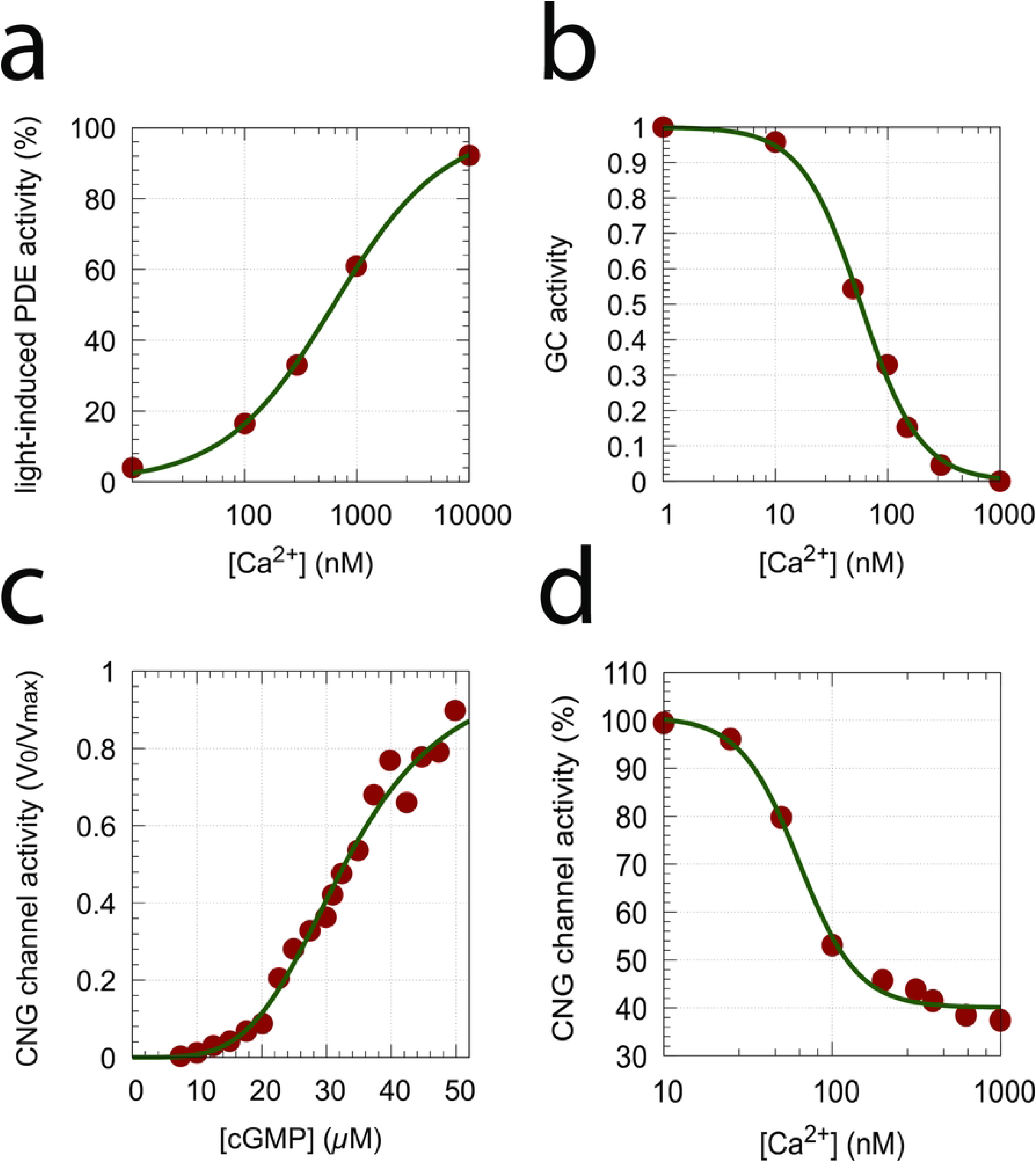
Experimental data used to extract parameter values. (a) Light-induced PDE activity as a function of Ca^2+^ concentration [35,36]; (b) Inhibition of GC activity by Ca^2+^ [37]; (c) CNG channel activity as a function of cGMP concentration [38]; (d) CNG channel activity as a function of Ca^2+^ concentration [38].

Also using salamander rods, Fig 14b shows the inhibition of GC activity by Ca^2+^ when using 0.5 mM GTP (Fig 13 in [37]). The function

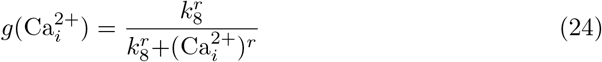

was fitted to the data with *k*_8_=(57.49±2.53)nM and *r*=1.65±0.12.

Using bovine retinae, Hsu and Molday [38] determined the influence of cGMP and Ca^2+^ on CNG channel activity in the presence of calmodulin (Fig 14, panels c and d, respectively). For CNG channel activation by cGMP (panel c) the following trial function

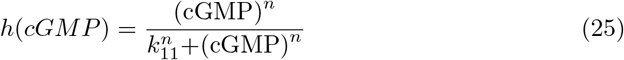

described the experimental data with *k*_11_=(32.81±0.39)*μ*M and *n*=4.14±0.23 quite well. For the inhibition of the CNG channel by Ca^2+^ (panel d) we fitted the function

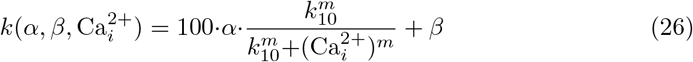

to the experimental data obtaining *α*=0.6067±0.0295, *k*_10_=(63.57±4.44)nM, *m*=2.50±0.38, and *β*=40.07±1.29. In Eq 21 *β*′ is given by *β*/100.

Organelles, such as mitochondria and the endoplasmatic reticulum (ER), store calcium with relative high concentrations (100-800*μ*M). There is evidence that intracellular Ca stores leak Ca into the cytosol [39–42]. Analyzing the data by Camello et al. [41] and Luik et al. [42], we observed (S4 Text and [43]) that the kinetics of the two recorded leaks were surprisingly different. While Camello et al. [41] found practically zero-order kinetics with respect to ER calcium and leak rates at around 0.25 *μ*M/s, the data by Luik et al. [42] show clean *first-order* kinetics with respect to ER calcium. Here Ca-dependent leak velocities between 5.5 and 0.36 *μ*M/s were observed (S4 Text). Also the results by Oldershaw et al. [39] and Missiaen et al. [40] indicate single or dual first-order kinetics in the decrease of store Ca. We wondered how calcium leaks may influence the photoadaptation of the model. As we will show in the section “Roles of the feedback loops” calcium leaks will have an influence on the steady state level of cGMP. In particular, when the leak rate 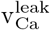 becomes larger than the K^+^ synthesis rate *k*_6_ in the NCKX-based calcium pump, then uncontrolled growth in 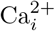 may occur (S4 Text).

cGMP hydrolysis in darkness (rate constant *k*_9_) is described as a first-order reaction with respect to cGMP. The value of *k*_9_ is taken from the modeling work by Nikonov et al. (Table IV in [44]) with *k*_9_=1.0s^-1^. The rates for the light-induced removal of cGMP (described by *k*_2_ and *k*_4_) are variable (light-dependent) parameters.

Parameter *k*_3_ represents the maximum rate of cGMP synthesis at low 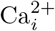 concentrations. Its value (*k*_3_=50 *μ*M/s) has been taken from the work by Nikonov et al. [44].

The extrusion of 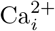 by NCKX is simplified as a second-order process with rate constant *k*_7_, i.e. 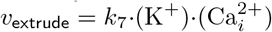. The influence of sodium ions on NCKX is not considered. It is interesting to note that in the absence of the CNG channel inhibition by 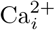 the NCKX pump would lead to robust perfect adaptation in cGMP by antithetic feedback [14], like the zero-order removal of *E* in the above idealized controllers (see for example, Eq 12). However, such an antithetic control of cGMP without CNG channel inhibition by 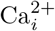 would lead to high and possibly toxic 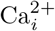 concentrations.

The remaining parameters *k*_5_, *k*_6_, and *k*_7_ have been chosen such that cGMP and 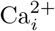 levels are close to observed experimental values, i.e. using *k*_5_=100 *μ*M/s, *k*_6_=0.5 *μ*M/s, and *k*_7_=2.0 *μ*M^-1^*s*^-1^. While *k*_7_ has no influence on the steady state values of cGMP and 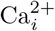 it has a significant influence on how fast steady state levels are approached after light perturbations are applied (S5 Text).

#### Application of pulse perturbations

In the majority of experiments on rod or cone cells light perturbations are applied in form of flashes in the millisecond range (see for example Fig 12). Fig 15 shows the application of 10 ms pulses of light in the model. A *k*_2_ pulse from 1 → 50 s^-1^ is applied at time *t*=1.0 s for different *k*_4_ backgrounds. In panel a the graphs are scaled such that the steady state levels of cGMP are set to zero and the individual excursions in cGMP can be compared. As for the above derepression controllers m2, m4, m6 and m8 the excursion ΔcGMP_max_ of the controlled variable cGMP decreases with increasing backgrounds while the speed of resetting to its original steady state increases with increasing backgrounds (Fig 15a). These changes are considered to be typical for the light adaptation in vertebrate photoreceptors (for example, see Ch. V in [31] and Fig 22-19C in [45]).

**Fig 15.**
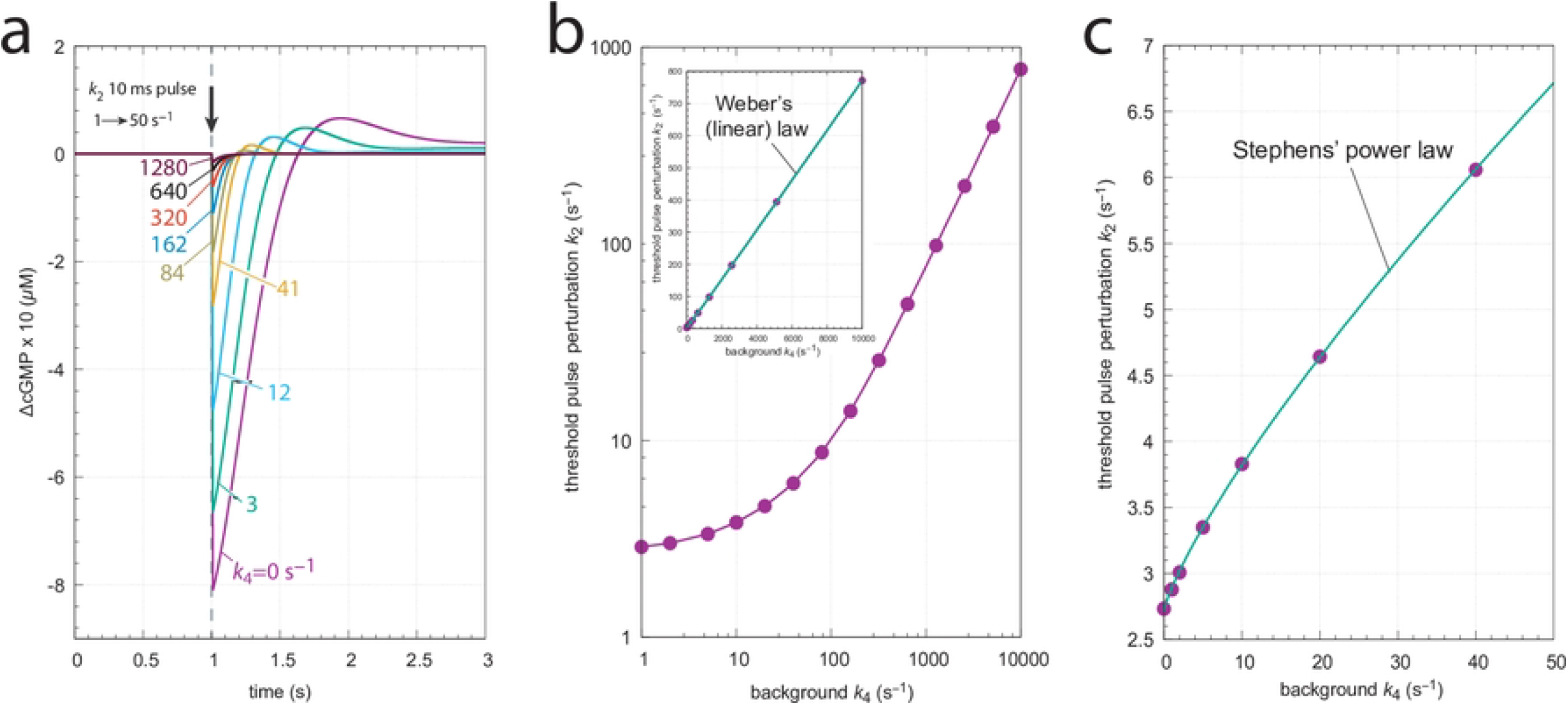
Application of 10 ms *k*_2_ pulses (1 →50 s^-1^) at different *k*_4_ backgrounds. (a) the scaled ΔcGMP levels against time. Colored numbers indicate the different background levels in s^-1^. Initial concentrations (in μM): cGMP_0_=9.04191, 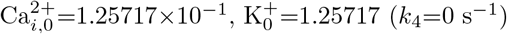 (*k*_4_=0 s^-1^); cGMP_0_=8.80039, 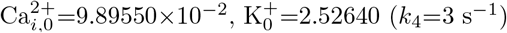 (*k*_4_=3 s^-1^); cGMP_0_=8.36375, 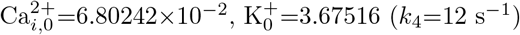 (*k*_4_=12 s^-1^); cGMP_0_=7.86039, 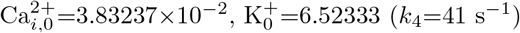 (*k*_4_=41 s^-1^); cGMP_0_=7.67946, 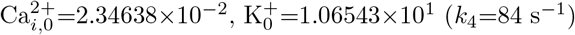 (*k*_4_=84 s^-1^); cGMP_0_=7.61322, 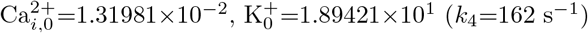 (*k*_4_=162 s^-1^); cGMP_0_=7.59537, 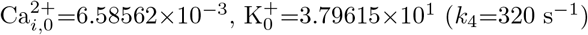 (*k*_4_=320 s^-1^); cGMP_0_=7.59210, 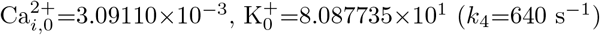 (*k*_4_=640 s^-1^); cGMP_0_=7.59160, 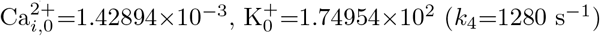 (*k*_4_=1280 s^-1^). Panel b shows the threshold perturbation *k*_2_, which leads to a ΔcGMP of 0.03 *μ*M as a function of background. The overall curved log-log plot turns out to be linear and follows Weber’s law (inset) as: threshold perturbation *k*_2_ = *a*·(*k*_4_)^*n*^+*b* with *a*=(0.069±0.001)*s*^*n*–1^, *n*=1.012±0.002, and *b*=(2.73±0.20)s^-1^. Panel c shows that at low backgrounds the threshold-background relationship follows Stephens’ power law, i.e., threshold perturbation *k*_2_ = *a*·(*k*_4_)^*n*^+*b* with *a*=(0.175±0.006)s^*n*–1^, *n*=0.800±0.009, and *b*=(2.72±0.01)s^-1^. Parameter and rate constant values are as described in the previous section. See also ‘S1 Programs’.

Fig 15b shows threshold light pulse (10 ms) perturbations *k*_2_ with a ΔcGMP of 0.03 *μ*M as a function of background light intensity *k*_4_. The main graph shows the log-log plot which resembles the experimental results with rods or cones (see Fig 22-19B in [45]). The inset shows that the threshold-background relationship is linear in agreement with Weber’s law, at least for large backgrounds. Panel c shows, on the other hand, that for small backgrounds the threshold-background relationship follows Stephens’ power law. In fact, replotting the original experimental data [46] shown in Fig 22-19B of Ref [45], indicates that Stephens’ law describes best the situation at low backgrounds, while at higher backgrounds the threshold-background relationship tends towards Weber’s law (S7 Text).

#### Application of step perturbations

We applied step perturbations in the model to see to what extent the CNG channel inhibition by calcium affects cGMP homeostasis and avoids robust perfect adaptation. Fig 16a shows the influence of *k*_2_ 1 → 50 s^-1^ steps at different backgrounds. The steps occur at time *t*=0.5 s and changes in cGMP are followed for 3 s. We also measured the maximum excursion of cGMP (ΔcGMP_max_) from its initial steady state level and the time *t_max_* at which ΔcGMP_max_ occurs (see inset).

**Fig 16.**
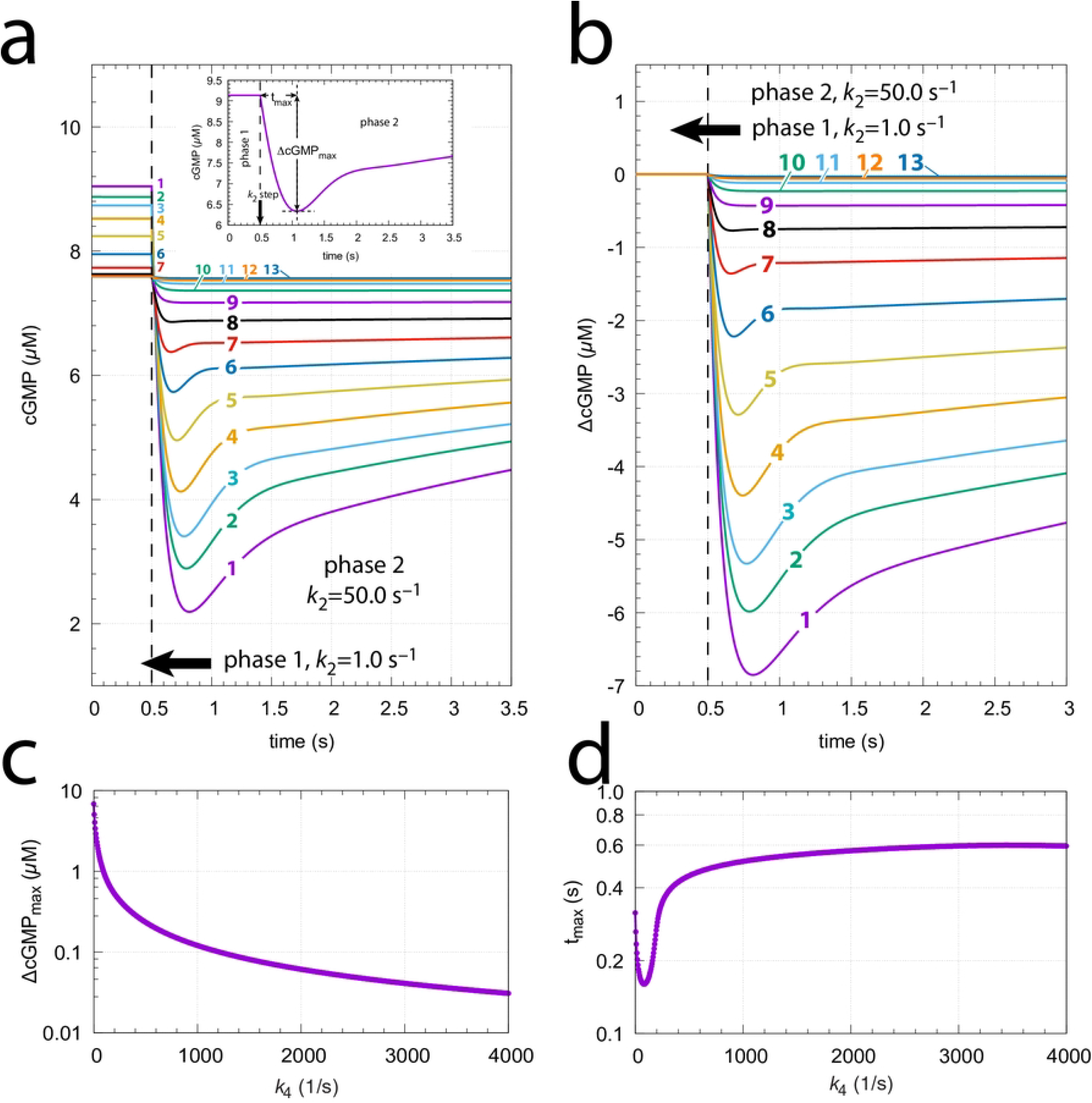
The model’s response towards *k*_2_ 1 →50 s ^−1^ steps at different backgrounds *k*_4_. (a) Unscaled cGMP concentrations as a function of time. The steps occur at *t*=0.5 s. Background *k*_4_ values (s^-1^): **1**, 0.0; **2**, 2.0; **3**, 4.0; **4**, 8.0; **5**, 16.0; **6**, 32.0; **7**, 64.0; **8**, 128.0; **9**, 256.0; **10**, 512.0; **11**, 1024.0; **12**, 2048.0; **13**, 4096.0. Initial concentrations (in *μ*M): cGMP_0_=9.04191, 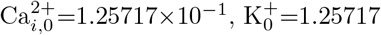 (*k*_4_=0 s^-1^); cGMP_0_=8.87243, 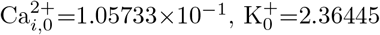 (*k*_4_=2 s^-1^); cGMP_0_=8.73490, 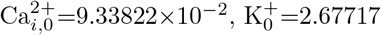 (*k*_4_=4 s^-1^); cGMP_0_=8.52196, 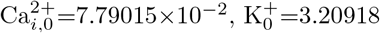 (*k*_4_=8 s^-1^); cGMP_0_=8.24168, 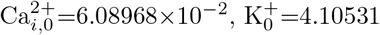 (*k*_4_=16 s^-1^); cGMP_0_=7.95044, 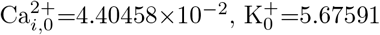 (*k*_4_=32 s^-1^); cGMP_0_=7.73313, 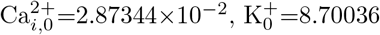 (*k*_4_=64 s^-1^); cGMP_0_=7.62877, 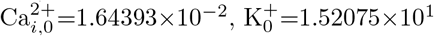 (*k*_4_=128 s^-1^); cGMP_0_=7.59845, 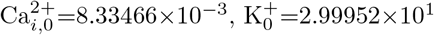 (*k*_4_=256 s^-1^); cGMP_0_=7.59259, 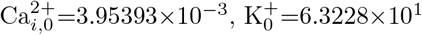 (*k*_4_=512 s^-1^); cGMP_0_=7.59165, 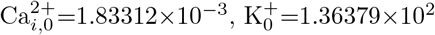 (*k*_4_=1024 s^-1^); cGMP_0_=7.59154, 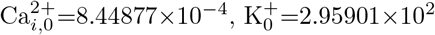 (*k*_4_=2048 s^-1^); cGMP_0_=7.59152, 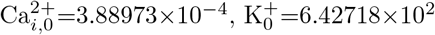 (*k*_4_=4096 s^-1^). Inset: Defining ΔcGMP_max_ and *t*_max_. (b) cGMP data as in (a), but scaled relative to their initial steady state concentrations. (c) and (d) ΔcGMP_max_ and *t*_max_ values as a function of backgrounds *k*_4_, respectively. Parameter and rate constant values are as described in section “Estimation of model parameters” (see also S1 Programs).

Fig 16b shows the same data as in (a), but scaled relative to their initial steady states. Due to the inhibition of CNG channels by calcium (Fig 13) the model does not show robust perfect adaptation (S5 Text, Fig 17). cGMP steady state levels during the step become significantly lower than their initial values before the step. This is seen in Fig 16a, where the pre-step steady state levels decrease as the background *k*_4_ increases. Not unexpected we see that with increasing backgrounds the ΔcGMP_*max*_ excursions decrease monotonically (Fig 16c). Surprisingly, however, we find that t_max_ first decreases, but then increases again (Fig 16d). Interestingly, when studying turtle photoreceptors, an increase of t_max_ at increasing backgrounds has also been reported by Baylor and Hodgkin [47]. They studied both flashes and steps [48, 49] and provided several models [50] to explain the lengthening of the peak time *t*_max_.

**Fig 17.**
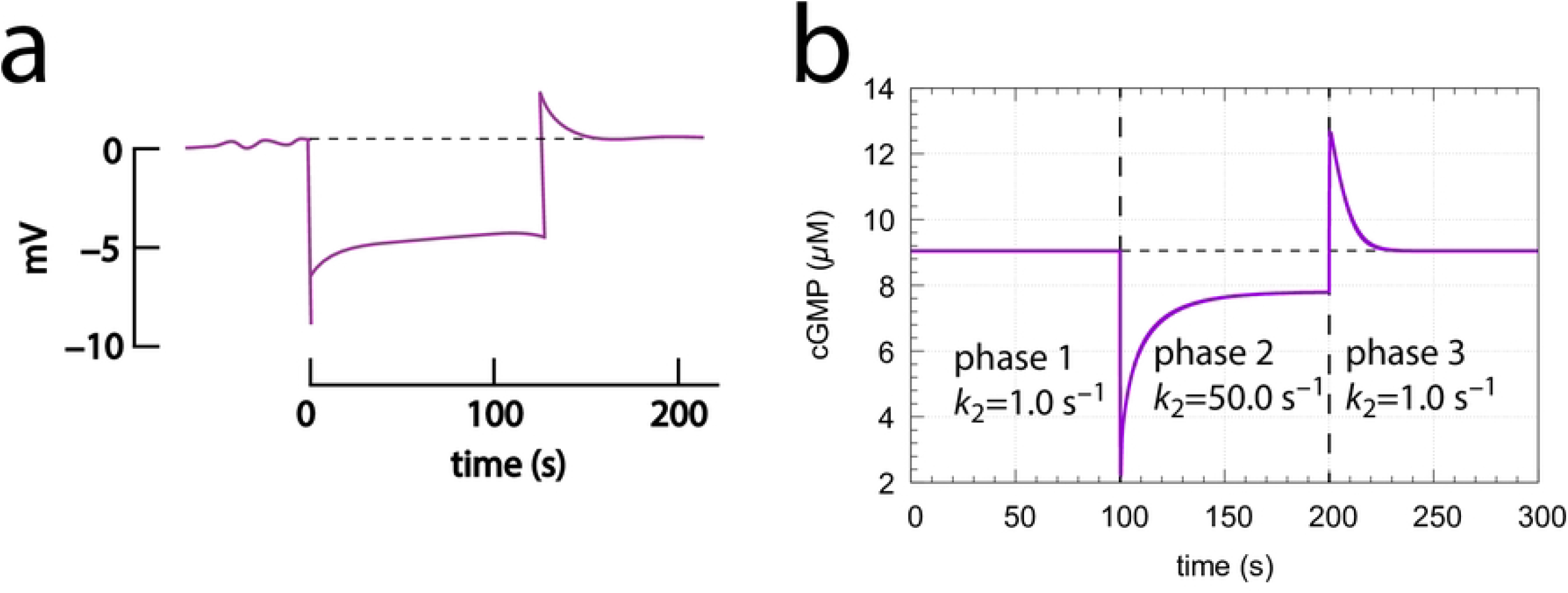
Experimental and model behaviors when applying step perturbations. (a) Experimental response of a red-sensitive turtle cone to a long step of light. Redrawn from Ref [47] (Fig 14, trace 2). (b) Model calculation using a *k*_2_ 1 → 50 s^-1^ step at time *t*=100s. After 100 s *k*_2_ returned to its original value. Background *k*_4_=0.0 s^-1^. All other rate parameters are as described in section “Estimation of model parameters”. Initial concentrations: cGMP=9.04*μ*M, 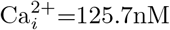, *K*^+^=2.0*μ*M. See also S1 Programs.

Fig 17a shows experimental results by Baylor and Hodgkin [47] when long steps of light are applied to red-sensitive turtle cones. The behavior of our model (panel b) is analogous with a typical overshooting when the step ends.

#### Roles of the feedback loops

Outlined in Fig 18 are the three feedback loops in the model. Feedback loops 1 and 2, both based on the inflow activation of 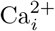 by cGMP (outlined in purple), feed respectively back to cGMP by a 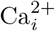-based inhibition (derepression) of cGMP synthesis (loop 1, analogous to m2, outlined in red) and by a 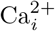-based activation of cGMP turnover (loop 2, analogous to m5, outlined in blue). Both loops 1 and 2 promote robust perfect cGMP homeostasis by antithetic control and oppose perturbations on cGMP. Feedback 3 (outlined in orange) keeps Ca 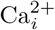 levels low to avoid high and cytotoxic calcium levels inside the cell.

**Fig 18.**
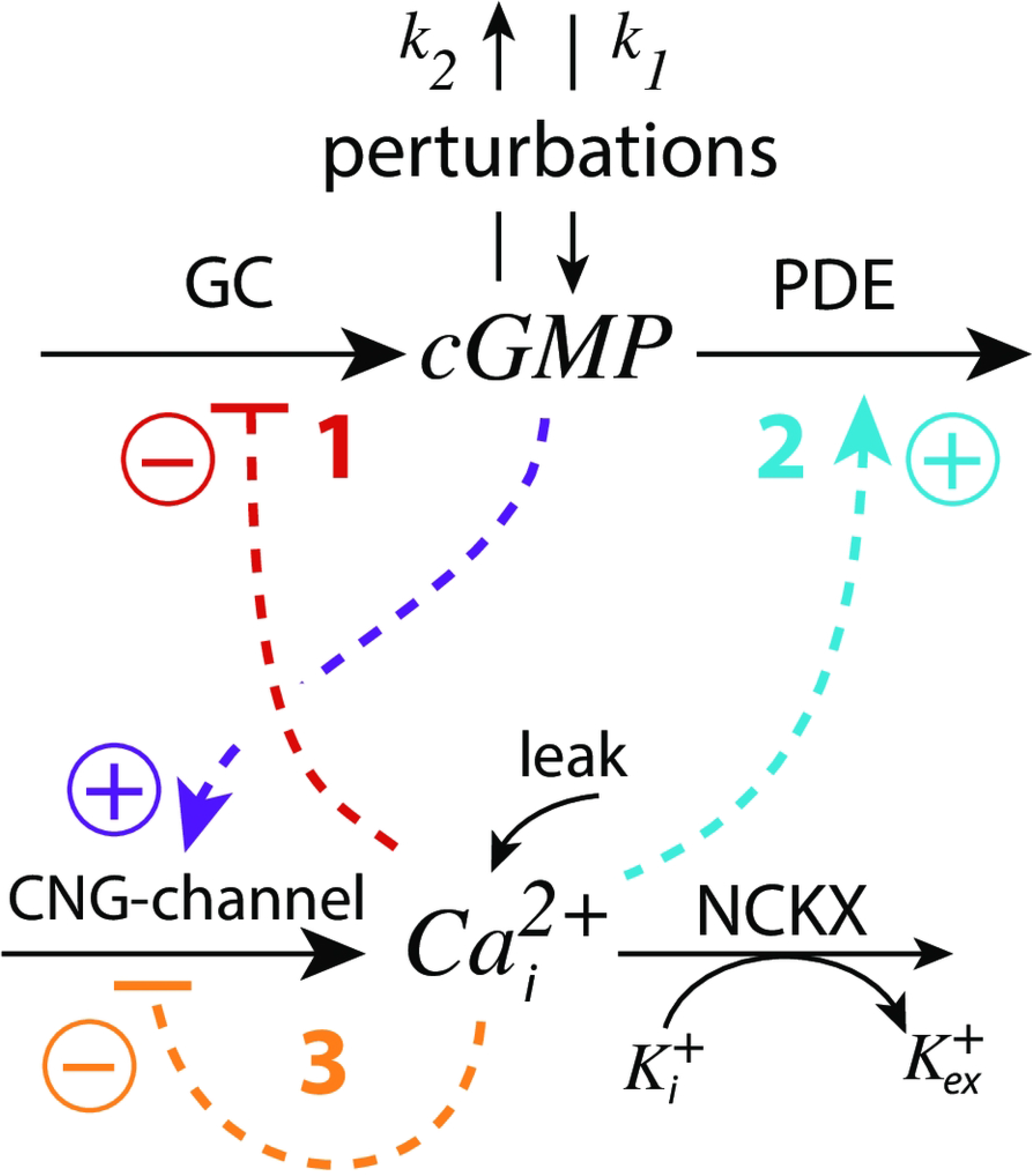
Schematic outline of the feedback loops 1-3 in the model (Fig 13). CNG: cyclic nucleotide-gated; GC: guanylate cyclase; PDE: phospho-diesterase; NCKX: potassium-dependent sodium-calcium exchangers (without the sodium part).

When feedback loop 3 is absent, for example by low 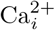 levels, the 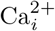-inhibition term in Eq 21 becomes 1, because

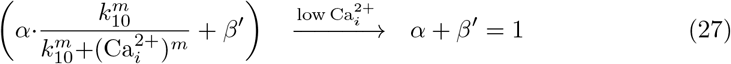

The remaining feedbacks 1 and 2 will provide robust perfect adaptation of cGMP, provided that there are sufficiently high GC and PDE activities to work as compensatory fluxes. This robust perfect adaptation in cGMP is due to the simultaneous NCKX-based removal of 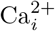 and K^+^ described by the term *k*_7_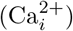(K^+^) in Eqs 21 and 22. The *k*_7_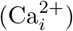(K^+^) transport term leads to robust antithetic integral control [14]. Instead of using *k*_7_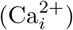(K^+^), one may have explicitly included the NCKX transporter protein catalyzing the move of 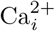 and K^+^ out of the cell under the dual-E control of 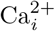 and K^+^ [16]. Anyway, using the *k*_7_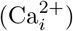(K^+^) term, the set-point of cGMP (*cGMP_set_*) is calculated by setting Eqs 21 and 22 to zero and solving for cGMP. The resulting steady state concentration of cGMP becomes cGMP’s set-point:

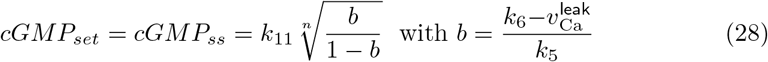

Using the experimentally determined rate parameters (see section “Estimation of model parameters”) leads to *cGMP_set_*=7.61*μ*M. The two feedback loops 1 and 2 act as an *antagonistic* pair as they will defend *cGMP_set_* robustly against both inflow and outflow perturbations, respectively. Fig 19a shows the homeostatic behavior of the loop 1-2 antagonistic feedback during three different phases where either inflow perturbation *k*_1_ or outflow perturbation *k*_2_ dominate. Although the antagonistic feedback can deal well with both inflow and outflow perturbations it needs sufficiently large GC and PDE activities, reflected by sufficiently high *k*_2_, *k*_3_, and *k*_4_ values, in order to provide the necessary compensatory fluxes.

**Fig 19.**
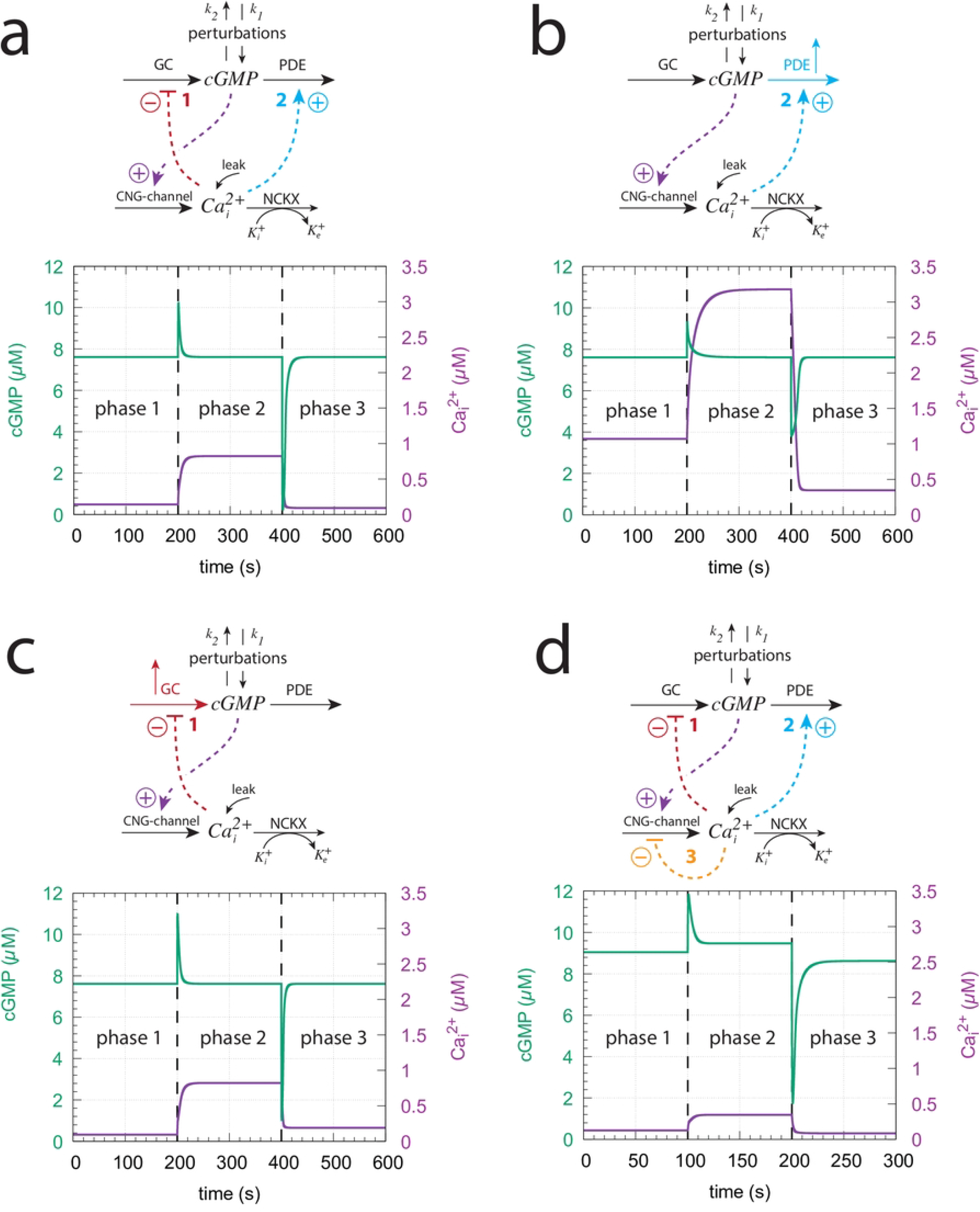
Influence of the model’s three feedback loops on the homeostatic behavior of cGMP and 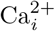. Perturbation profile in panels (a)-(d): phase 1: *k*_1_=0.0*μ*M/s, *k*_2_=1.0s^-1^, *k*_4_=0.0s^-1^; phase 2: *k*_1_=7.0*μ*M/s, *k*_2_=0.0s^-1^, *k*_4_=0.0s^-1^; phase 3: *k*_1_=0.0*μ*M/s, *k*_2_=7.0s^-1^, *k*_4_=0.0s^-1^. (a) Both feedback 1 and 2 are operative. Robust homeostasis of cGMP is observed with *cGMP_set_*=7.61*μ*M. Other rate constants values are as described in section “Estimation of model parameters”. Initial concentrations: cGMP=7.612*μ*M, 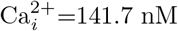, K^+^=1.760*μ*M. (b) Feedback 2 is only operative. In order to keep cGMP at its set-point *k*_4_ needs to be increased to 8.0 *μ*M/s in all three phases (indicated in the scheme by the blue upright arrow). Initial concentrations: cGMP=7.612*μ*M, 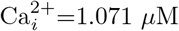 *μ*M, K^+^=2.335*μ*M. (c) Feedback 1 is only operative. To keep cGMP at its set-point *k*_3_ has been increased from 50.0 *μ*M/s to 500.0 *μ*M/s in phase 3 (indicated in the scheme by the red upright arrow). Initial concentrations: cGMP=7.612*μ*M, 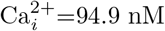, K^+^=2.64*μ*M. (d) All feedback loops are operative with rate constants as in panel (a). Although perfect adaptation in cGMP is lost both cGMP and 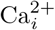 undergo only small variations when the perturbations are applied with lowest 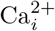 levels. Initial concentrations: cGMP=9.042*μ*M, 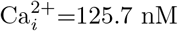, K^+^=1.989*μ*M. See S4 Text how the leak term affects this configuration.

Fig 19b shows the system’s behavior when only feedback loop 2 is operative. To achieve control by only feedback 2 the condition in Eq 27 needs to hold and the inhibition of GC by 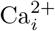 has to be abolished by using a high inhibition constant *k*_8_. We have used *k*_8_=1×10^9^*μ*M with r=1.0. When applying the same perturbation profile as in Fig 19a it turned out that the PDE activity from Fig 19a was not sufficient to keep cGMP homeostasis at *cGMP_set_*=7.61*μ*M. The reason for this is that the lack of feedback loop 1 causes a higher cGMP and Ca^2+^ inflow into the cell. When becoming too high the Ca^2+^ inflow cannot be absorbed by the constant 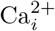 removal speed *k*_6_ of NCKX. In other words, the antithetic zero-order removal kinetics of 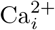 by NCKX will become too slow and thereby lead to a steady increase (windup) in the concentration of 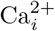 (S5 Text). To avoid this and to keep cGMP robustly at *cGMP_set_*=7.61*μ*M we have in Fig 19b increased the background *k*_4_ to 8 *μ*M/s (indicated by the blue upright arrow). Alternatively, one may increase the constant removal speed *k*_6_ of the NCKX pump, but this will result in a change of *cGMP_set_* (see also S6 Text).

Fig 19c shows the system’s behavior when only feedback loop 1 is present. To get only loop 1 operative the condition of Eq 27 is imposted and the activation constant *k*_12_ (Fig 13) is set to zero. To act as a robust inflow controller cGMP homeostasis requires sufficiently high *k*_3_ values. With the perturbation profile from panel (a) *k*_3_ needs to be increased in phase 3 by one order of magnitude to *k*_3_=500*μ*/s (indicated by the red upright arrow in Fig 19c) in order to avoid cGMP levels below *cGMP_set_*=7.61*μ*M (see also S6 Text).

When all three loops are operative (Fig 19d) the robust perfect adaptation of cGMP is lost due to the presence of feedback loop 3. However, with respect to the applied perturbations cGMP levels show only small variations and 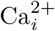 steady state concentrations have their lowest values. The results in Fig 19 show that the antagonistic feedback between loops 1 and 2 is more efficient than when loops 1 or 2 are isolated. Although the robust perfect adaptation of cGMP is lost in the presence of feedback loop 3, the overlayed feedback structure between all three feedbacks provides a compromise between robust perfect adaptation of cGMP and the need to avoid high cytotoxic 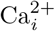 levels.

Another aspect of the three feedbacks’ overlay concerns the resetting times at varying/increasing backgrounds. While a faster resetting with increasing backgrounds has been described as a typical property of vertebrate photoadaptation (see section V in [31]), in turtle photoreceptors Baylor et al. [47] found that increasing backgrounds first lead to a decrease in peak time (analogous to *t_max_*), but further increases of the background eventually lead to an increase of the peak time (*t_max_*), as qualitatively observed in Fig 16d. The increase of the time to peak was explained by Baylor et al. [50] by a hypothetical autocatalytic reaction which removed particles blocking the ionic channels. An additional factor could be a differential dominance between feedback loops 1 and 2, since loop 1 and loop 2 affect the resetting differently analogous as described for the m2 (Fig 8) and m5 (S1 Text) controllers.

Fig 20 shows ΔcGMP and *t_max_* as a function of the feedback arrangement. In panel (a) we have feedback loops 1 and 3 combined, while in panel (b) we have only feedback loop 2. When testing a 1.0 →50.0 *μ*M/s *k*_2_ step for increasing backgrounds both feedback arrangements show a monotonic decline of ΔcGMP as a function of background *k*_4_ (middle panels), but differ in their *t_max_* responses (bottom panels). While combined feedback loop 1 and 3 show a monotonic shortening of *t_max_*, in the feedback 2 arrangement *t_max_* first decreases, but then increases again as background *k*_4_ increases, as found experimentally by Baylor et al. [47] and when all three feedback loops are combined (Figs 16c and d). Since the single feedback 2 behavior (Fig 20 b) resembles that of all three feedbacks combined (Figs 19c and d) we conclude that in our model with the used parameter values feedback 2 is dominating over the two other feedbacks with respect to the system’s resetting behavior. In organisms where the photoadaptation shows faster resettings (decreasing or constant *t_max_*) with increasing backgrounds, as found in Ref. [30] and highlighted in the review by Fain et al. [31], the feedback loop 2 may be weakened and loops 1 and 3 may become more dominant. Since the rate parameters of our model were taken from different organisms it is possible that these combined parameters reflect a situation closer to turtles [47] than, for example, to Macaque monkeys [30].

**Fig 20.**
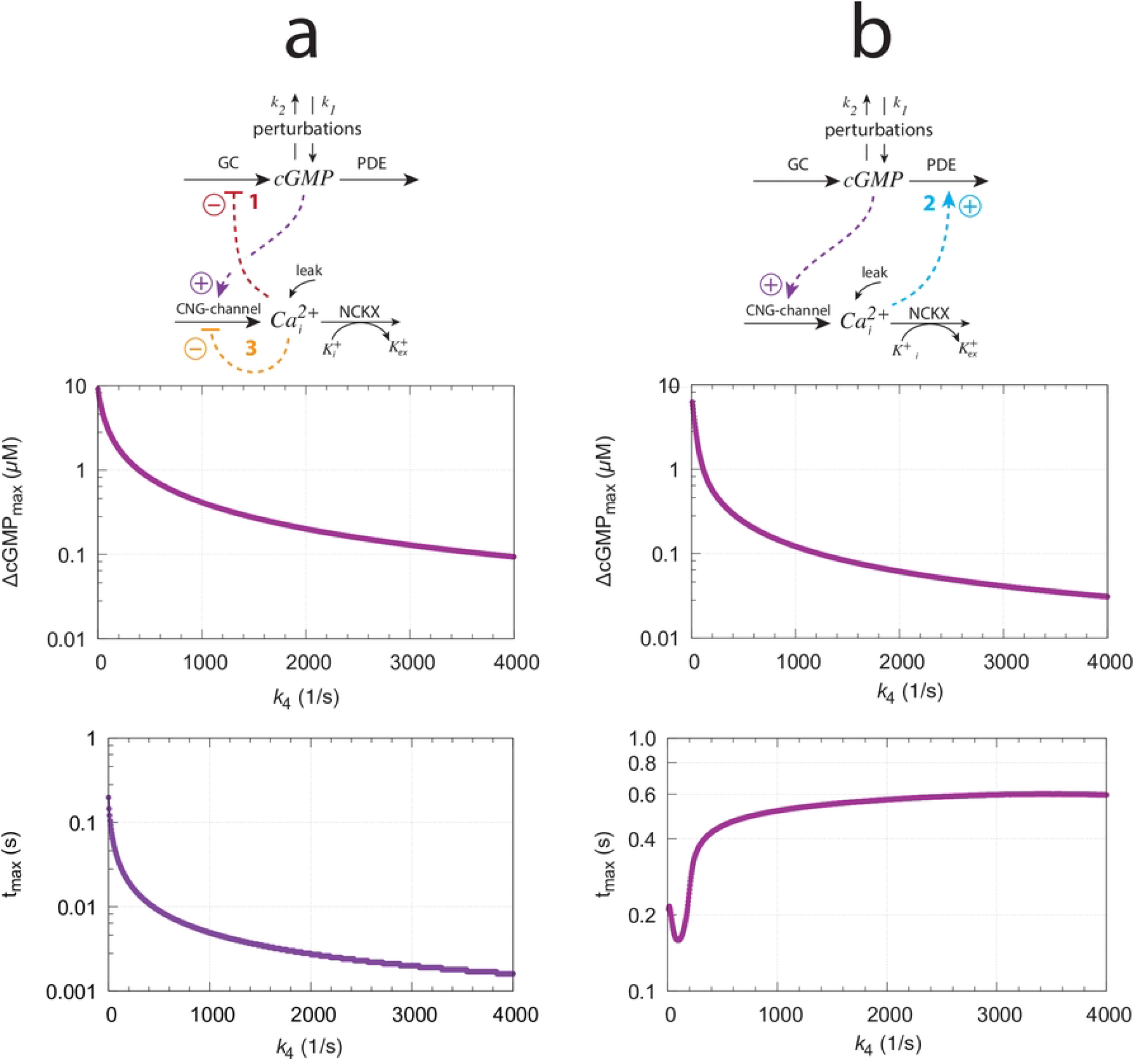
The model’s resetting behavior for different feedback arrangements when applying a 1.0 →50.0 s^-1^ step in *k*_2_ as a function of backgrounds *k*_4_. (a) Feedback loops 1 and 3 are combined. (b) Feedback 2 only. Used parameter values, rate constants, and definition of ΔcGMP and *t_max_* are as in Fig 16.

## Conclusion and outlook

Studying perturbations with backgrounds on eight basic feedback loops m1-m8 with integral control show that these homeostatic controllers divide into two classes dependent on how the compensatory flux is activated. In the class where the compensatory flux is based on derepression faster resetting with respect to a standard step perturbation is observed when backgrounds increase. In the other class when compensatory fluxes are based on direct activation the resetting to the set-point slows down as backgrounds increase. In both cases the maximum excursion of the controlled variable following the perturbation decrease monotonically as backgrounds increase. We originally thought that vertebrate photoadaptation would be a nice example of using sole derepression kinetics in a robust control of cGMP with cellular calcium as the controller. However, the situations turned out to be more complex with an overlay of three feedback loops, one based on derepression by Ca^2+^ on GC (feedback 1) and one based on Ca^2+^-based light activation of PDE (feedback 2). The antagonistic pair of combined feedbacks 1 and 2 show more improved properties than each of the individual controllers alone. In addition, there is a third Ca^2+^-controlling feedback (feedback 3) which apparently avoids high cytotoxic Ca^2+^ levels. This combination of three feedback loops indicates that robust perfect adaptation of cGMP by feedback loops 1 and 2 is not by itself an evolutionary target, but that a compromise between these three controllers has developed by keeping both cGMP *and* cytosolic Ca^2+^ levels at narrow limits, but not by robust perfect adaptation mechanisms. Furthermore, there is also evidence that photoadaptation with increasing backgrounds may both accelerate or slow down the resetting kinetics dependent on the dominance of feedback 1 or feedback 2.

The findings that controllers m1-m8 react so differently on perturbations with respect to backgrounds may be of importance also in other physiological systems. For example, blood sugar levels are controlled by two major feedback loops involving insulin and glucagon. Since glucose control by insulin is based by an activation of beta cells via glucose (see Supporting Material in Ref. [11]), constantly high glucose levels (“glucose overload”) [51,52], for example, may lead to a slower resetting of the insulin-based control loop in comparison with more rapid anticipated adaptations at lower glucose levels. Such a slowing-down response may be one of the causes that could participate in the mechanisms leading to insulin resistance and early diabetes. To what extent these aspects of background perturbations in homeostatic systems apply to the development of diabetes or have implications in other homeostatic systems needs certainly further investigations.

## Supporting information

**S1 Programs. Documentation.** A zip-file with python scripts describing the results for motifs m1 (Fig 4a), m7 (Fig 6a), m2 (Fig 8a), m8 (Fig 11a), m3 and m5 (S1 Text, Figs S2a and S4a), m4 (S2 Text, Fig S2), m6 (S2 Text, Fig S4a), Fig 15, Fig 16a,b, Fig 17b, and Fig 19.

**S1 Text. Response kinetics of controllers m3 and m5.** Applied step perturbations lead to slower resetting kinetics for increasing backgrounds.

**S2 Text. Response kinetics of controllers m4 and m6.** Applied step perturbations lead to faster resetting kinetics for increasing backgrounds.

**S3 Text. Response kinetics controller m2 with antithetic integral control.** The behavior is dynamically identical to that of m2 with zero-order kinetics.

**S4 Text. Influence of Ca leak kinetics on photoadaptation.** A comparison how experimentally observed zero-order and first-order Ca leak kinetics affect photoadaptation in the model and when homeostatic breakdown occurs.

**S5 Text. Influence of *k*_5_, *k*_6_, and *k*_7_ on the model’s photoadaptation.** By using a *k*_1_-*k*_2_ perturbation profile influences of *k*_5_, *k*_6_, and *k*_7_ on the model’s resetting kinetics are shown.

**S6 Text. Experimental light adaptation data.** Replots of experimental data show, as indicated by model calculations, that Stephens’ law is followed at low backgrounds, while at higher backgrounds the response tends towards Weber’s law.

## References

1. Cannon W. Organization for Physiological Homeostatics. Physiol Rev. 1929;9:399–431.

2. Langley, LL, editor. Homeostasis. Origins of the Concept. Stroudsbourg, Pennsylvania: Dowden, Hutchinson & Ross, Inc.; 1973.

3. Mrosovsky N. Rheostasis. The Physiology of Change. New York: Oxford University Press; 1990.

4. Sterling P, Eyer J. Allostasis: A new paradigm to explain arousal pathology. In: Fisher, S and Reason, J, editor. Handbook of Life Stress, Cognition and Health. New York: John Wiley & Sons; 1988. p. 629–49.

5. Schulkin J. Rethinking Homeostasis. Allostatic Regulation in Physiology and Pathophysiology. Cambridge, Massachusetts: MIT Press; 2003.

6. Schulkin J. Allostasis, Homeostasis and the Costs of Physiological Adaptation. Cambridge, Massachusetts: Cambridge University Press; 2004.

7. Moore-Ede M. Physiology of the circadian timing system: Predictive versus reactive homeostasis. Am J Physiol. 1986;250:R737–52.

8. Lloyd D, Aon M, Cortassa S. Why Homeodynamics, Not Homeostasis? The Scientific World. 2001;1:133–145.

9. Carpenter R. Homeostasis: A plea for a unified approach. Advances in Physiology Education. 2004;28(4):180–187.

10. Ruoff P, Nishiyama N. Frequency switching between oscillatory homeostats and the regulation of p53. PloS One. 2020;15(5):e0227786.

11. Drengstig T, Jolma IW, Ni XY, Thorsen K, Xu XM, Ruoff P. A Basic Set of Homeostatic Controller Motifs. Biophys J. 2012;103:2000–2010.

12. Wilkie J, Johnson M, Reza K. Control Engineering. An Introductory Course. New York: Palgrave; 2002.

13. Ni XY, Drengstig T, Ruoff P. The control of the controller: Molecular mechanisms for robust perfect adaptation and temperature compensation. Biophys J. 2009;97:1244–53.

14. Briat C, Gupta A, Khammash M. Antithetic integral feedback ensures robust perfect adaptation in noisy biomolecular networks. Cell Systems. 2016;2(1):15–26.

15. Aoki SK, Lillacci G, Gupta A, Baumschlager A, Schweingruber D, Khammash M. A universal biomolecular integral feedback controller for robust perfect adaptation. Nature. 2019;570(7762):533–537.

16. Waheed Q, Zhou H, Ruoff P. Kinetics and mechanisms of catalyzed dual-E (antithetic) controllers. PloS One. 2022;17(8):e0262371.

17. Shoval O, Goentoro L, Hart Y, Mayo A, Sontag E, Alon U. Fold-change detection and scalar symmetry of sensory input fields. PNAS. 2010; p. 201002352.

18. Drengstig T, Ni XY, Thorsen K, Jolma IW, Ruoff P. Robust Adaptation and Homeostasis by Autocatalysis. J Phys Chem B. 2012;116:5355–5363.

19. Briat C, Zechner C, Khammash M. Design of a synthetic integral feedback circuit: dynamic analysis and DNA implementation. ACS Synthetic Biology. 2016;5(10):1108–1116.

20. Eisler H, Eisler AD, Hellström A. Psychophysical Issues in the Study of Time Perception. In: Grondin S, editor. Psychology of Time. Emerald Group Publishing Limited; 2008. p. 75–76.

21. Ross HE, Murray DJ. EH Weber on the Tactile Senses. Psychology Press; 2018.

22. Fechner GT. Elemente der Psychophysik. Zweiter Theil. Breitkopf und Härtel; 1860.

23. Stevens SS. On the psychophysical law. Psychological Review. 1957;64(3):153.

24. MacKay DM. Psychophysics of perceived intensity: A theoretical basis for Fechner’s and Stevens’ laws. Science. 1963;139(3560):1213–1216.

25. Radhakrishnan K, Hindmarsh AC. Description and Use of LSODE, the Livermore Solver for Ordinary Differential Equations. NASA Reference Publication 1327, Lawrence Livermore National Laboratory Report UCRL-ID-113855. Cleveland, OH 44135-3191: National Aeronautics and Space Administration, Lewis Research Center; 1993.

26. Warwick K. An Introduction to Control Systems. Second Edition. World Scientific; 2019.

27. Fjeld G, Thorsen K, Drengstig T, Ruoff P. Performance of homeostatic controller motifs dealing with perturbations of rapid growth and depletion. The Journal of Physical Chemistry B. 2017;121(25):6097–6107.

28. Drobac G, Waheed Q, Heidari B, Ruoff P. An amplified derepression controller with multisite inhibition and positive feedback. PLoS One. 2021;16(3):e0241654.

29. Drengstig T, Jolma I, Ni X, Thorsen K, Xu X, Ruoff P. A basic set of homeostatic controller motifs. Biophys J. 2012;103(9):2000–2010.

30. Schneeweis D, Schnapf J. Noise and light adaptation in rods of the macaque monkey. Visual Neuroscience. 2000;17(5):659–666.

31. Fain GL, Matthews HR, Cornwall MC, Koutalos Y. Adaptation in vertebrate photoreceptors. Physiological Reviews. 2001;81(1):117–151.

32. Baylor DA, Nunn B, Schnapf J. The photocurrent, noise and spectral sensitivity of rods of the monkey Macaca fascicularis. The Journal of Physiology. 1984;357(1):575–607.

33. Krizaj D, Copenhagen DR. Calcium regulation in photoreceptors. Frontiers in Bioscience - Landmark. 2002;7(4):d2023–d2044.

34. Marks F, Klingmüller U, Müller-Decker K. Cellular Signal Processing: An Introduction to the Molecular Mechanisms of Signal Transduction. Second Edition. Garland Science; 2017.

35. Koutalos Y, Nakatani K, Yau K. The cGMP-phosphodiesterase and its contribution to sensitivity regulation in retinal rods. The Journal of General Physiology. 1995;106(5):891–921.

36. Koutalos Y, Yau KW. Regulation of sensitivity in vertebrate rod photoreceptors by calcium. Trends in Neurosciences. 1996;19(2):73–81.

37. Koutalos Y, Nakatani K, Tamura T, Yau K. Characterization of guanylate cyclase activity in single retinal rod outer segments. The Journal of General Physiology. 1995;106(5):863–890.

38. Hsu YT, Molday RS. Modulation of the cGMP-gated channel of rod photoreceptor cells by calmodulin. Nature. 1993;361(6407):76–79.

39. Oldershaw KA, Nunn D, Taylor CW. Quantal Ca^2+^mobilization stimulated by inositol 1, 4, 5-trisphosphate in permeabilized hepatocytes. Biochemical Journal. 1991;278(3):705–708.

40. Missiaen L, Smedt HD, Parys JB, Raeymaekers L, Droogmans G, Bosch LVD, et al. Kinetics of the non-specific calcium leak from non-mitochondrial calcium stores in permeabilized A7r5 cells. Biochemical Journal. 1996;317(3):849–853.

41. Camello C, Lomax R, Petersen OH, Tepikin A. Calcium leak from intracellular stores - the enigma of calcium signalling. Cell Calcium. 2002;32(5-6):355–361.

42. Luik RM, Wang B, Prakriya M, Wu MM, Lewis RS. Oligomerization of STIM1 couples ER calcium depletion to CRAC channel activation. Nature. 2008;454(7203):538–542.

43. Selstø CH, Ruoff P. A basic model of calcium homeostasis in non-excitable cells. bioRxiv. 2022; doi:doi.org/10.1101/2022.x.y.

44. Nikonov S, Lamb T, Pugh Jr EN. The role of steady phosphodiesterase activity in the kinetics and sensitivity of the light-adapted salamander rod photoresponse. The Journal of General Physiology. 2000;116(6):795–824.

45. Kandel ER, Koester JD, Mack SH, Siegelbaum S, editors. Principles of Neural Science. Sixth Edition. McGraw-Hill; 2021.

46. Wyszecki G, Stiles WS. Color Science. Second edition. Wiley New York; 1982.

47. Baylor D, Hodgkin A. Changes in time scale and sensitivity in turtle photoreceptors. The Journal of Physiology. 1974;242(3):729–758.

48. Baylor D, Hodgkin A. Detection and resolution of visual stimuli by turtle photoreceptors. The Journal of Physiology. 1973;234(1):163–198.

49. Baylor D, Hodgkin A, Lamb T. The electrical response of turtle cones to flashes and steps of light. The Journal of Physiology. 1974;242(3):685–727.

50. Baylor D, Hodgkin A, Lamb T. Reconstruction of the electrical responses of turtle cones to flashes and steps of light. The Journal of Physiology. 1974;242(3):759–791.

51. Moreira PI. High-sugar diets, type 2 diabetes and Alzheimer’s disease. Current Opinion in Clinical Nutrition & Metabolic Care. 2013;16(4):440–445.

52. Joubert M, Manrique A, Cariou B, Prieur X. Diabetes-related cardiomyopathy: The sweet story of glucose overload from epidemiology to cellular pathways. Diabetes & Metabolism. 2019;45(3):238–247.

